# Correction of Niemann-Pick type C1 disease with the histone deacetylase inhibitor valproic acid

**DOI:** 10.1101/724187

**Authors:** Kanagaraj Subramanian, Darren M Hutt, Vijay Gupta, Shu Mao, William E. Balch

## Abstract

Niemann-Pick type C (NPC) disease is primarily caused by mutations in the *NPC1* gene and is characterized by the accumulation of unesterified cholesterol and lipids in the late endosomal (LE) and lysosomal (Ly) compartments. The most prevalent disease-linked mutation is the I1061T variant of NPC1, which exhibits defective folding and trafficking from the endoplasmic reticulum to the LE/Ly compartments. We now show that the FDA-approved histone deacetylase inhibitor (HDACi) valproic acid (VPA) corrects the folding and trafficking defect associated with I1061T-NPC1 leading to restoration of cholesterol homeostasis, an effect that is largely driven by a reduction in HDAC7 expression. The VPA-mediated trafficking correction is in part associated with an increase in the acetylation of lysine residues in the cysteine-rich domain of NPC1. The HDACi-mediated correction is synergistically improved by combining it with the FDA-approved anti-malarial, chloroquine, a known lysosomotropic compound, which improved the stability of the LE/Ly-localized fraction of the I1061T variant. We posit that combining the activity of VPA, to modulate epigentically the cellular acetylome, with chloroquine, to alter the lysosomal environment to favor stability of the trafficked I1061T variant protein, can have a significant therapeutic benefit in patients carrying at least one copy of the I1061T variant of NPC1, the most common disease-associated mutation leading to NPC disease. Given its ability to cross the blood brain barrier, we posit VPA provides a potential mechanism to improve the response to 2-hydroxypropyl-β-cyclodextrin, by restoring functional NPC1 to cholesterol managing compartment as an adjunct therapy.

## Introduction

Understanding the impact of genetic diversity and its therapeutic management in the clinic is the forefront of precision medicine (1). Niemann-Pick type C disease (NPC) is an autosomal recessive, neurodegenerative disorder that arises in response to mutations in the *NPC1* and *NPC2* genes with the former accounting for 95% of cases affecting cholesterol homeostasis (CH) in the late <u>e</u>ndosome (LE) and <u>ly</u>sosome (Ly) compartments (LE/Ly)(2-5). NPC1 is a multi-membrane spanning 1278 amino acid protein with functional domains largely oriented towards the lumen of the Ly (**Fig. 1A**)(6-8). NPC1 is synthesized in the endoplasmic reticulum (ER) and transported to the LE/Ly compartments (9-12) where it works in tandem with NPC2 to ensure cholesterol distribution between all cellular and subcellular membranes (13-15). Like most rare diseases, genetic variation in the clinic contributes differentially to disease onset and progression (1,16). We have shown that variants triggering NPC1 disease are differentially responsive to both proteostasis regulators (17) and the HDAC inhibitor, SAHA (18), suggesting a potential route to manipulate the variant challenged NPC1 misfolding by stabilizing its fold.

**Figure 1.**
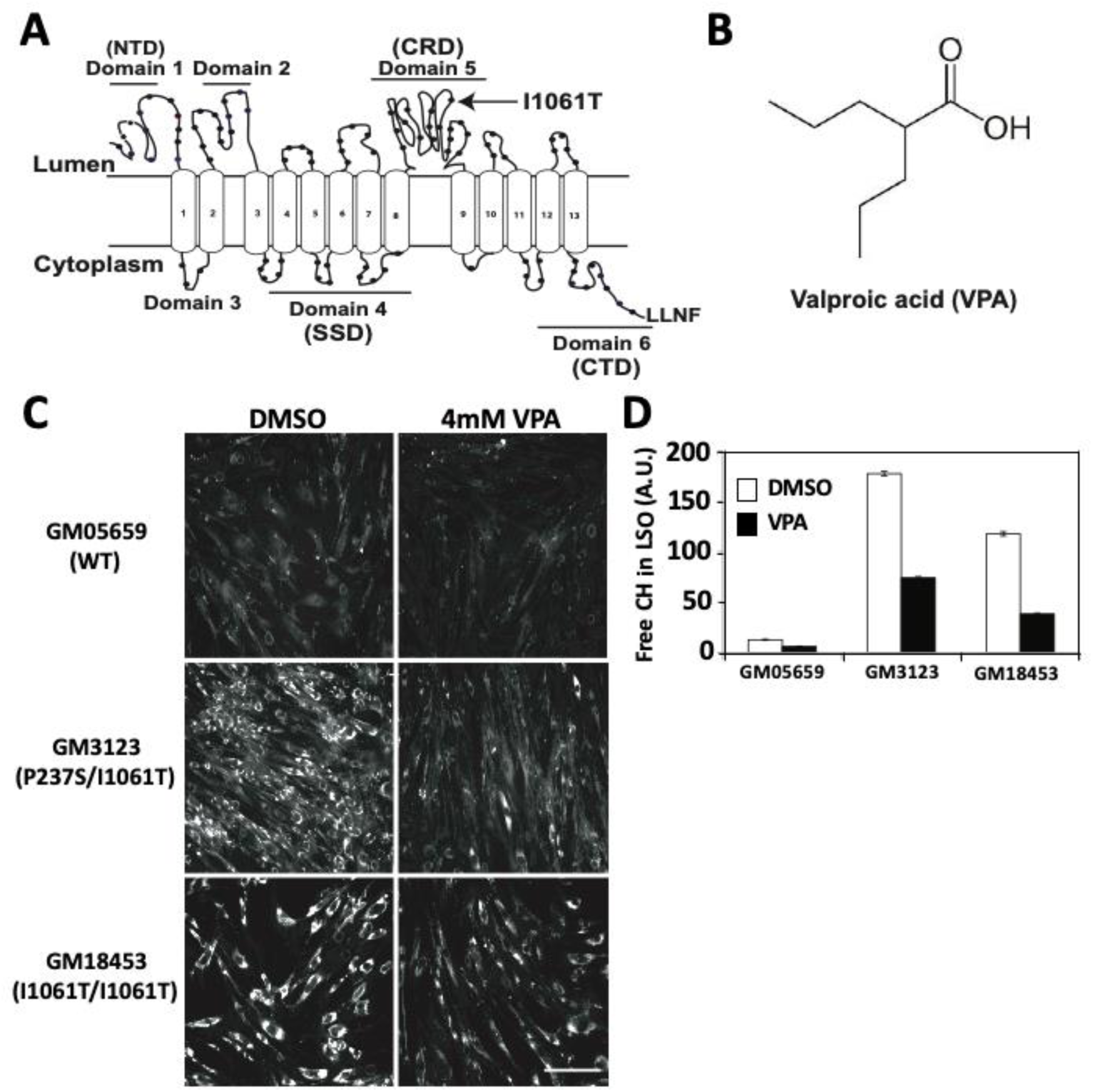
VPA reduces cholesterol accumulation in patient derived NPC1 fibroblasts. **A.** Schematic representation of the topology of human NPC1 protein and the location of the most prevalent I1061T mutant. The domain arrangement of NPC1 includes an N-terminal domain (NTD) (Domain 1), a sterol sensing domain (SSD) (domain 4), a Cysteine rich domain (CRD) (domain 5) and a C-terminal domain (CTD) (domain 6). **B.** Chemical structure of Valproic acid (VPA). **C.** Epifluorescence microscopy of filipin labeled free cholesterol (CH) in human fibroblasts expressing WT-NPC1 (GM05659), P237S/I1061T-NPC1 (GM3123) and I1061T/I1061T (GM18453) following the treatment with 4 mM VPA or vehicle for 48 h. Cells were imaged at a magnification of 10X and the scale bar represents 25 μm. **D.** Quantitative analysis of the filipin labeled free CH shown in panel **C**. The data represents the average ± SD of free CH in the Lysosomal storage organelle (LSO) (n = 3).

NPC1 disease progression is primarily a consequence of neuronal dysfunction in the hippocampus (19-23), The most common disease-associated mutation leading to NPC is the I1061T-NPC1, which accounts for 15-20% of all clinical cases (24,25). Disease presentation is characterized by the aberrant accumulation of unesterified cholesterol (CH), glycosphingolipids (GSL), sphingomyelin and sphingosine in the LE/Ly compartments (26) resulting in either a toxic accumulation of CH in the LE/Ly compartment or depletion of accessible CH by other cellular compartments (27) culminating in the progressive loss of Purkinje (PK) cells in the cerebellum. The loss of PK neurons causes ataxia, dysarthria, vertical supranuclear gaze palsy and a decline of neurological functions (27,28), phenotypic hallmarks of NPC1 disease.

More than 252 disease-causing mutations in *NPC1* have been reported in the clinic (29,30). These variants exhibit a distribution across the polypeptide sequence including variants in cytosolic, lumenal and transmembrane domains suggestive of a broadly metastable protein (31,32). Patient fibroblasts homozygous for the I1061T variant exhibit reduced protein expression and defective folding of NPC1, leading to its retention in the ER where it is subsequently degraded by the ubiquitin-proteasome system (UPS) (33). In contrast, other variants show efficient trafficking to the LE/Ly compartments but lack activity (17,18). Given the critical role played by NPC1 in cholesterol homeostasis, uncovering small molecules or biological pathways that restore the trafficking of a functional form of the I1061T variant to LE/Ly compartments will be critical for the treatment of NPC disease.

Current therapeutic opportunities for NPC disease include a clinical trial for the intrathecal administration of the cholesterol homeostasis modulator, 2-hydroxypropyl-β-cyclodextrin (HPβCD) (https://clinicaltrials.gov/ct2/show/NCT03879655), marketed as VTS-270, and arimoclomol, a heat shock protein (Hsp) activator (34-37). While, HPβCD has been shown to correct cholesterol homeostasis, behavioral and physiological symptoms in both mouse (38-40) and cat (41) models of disease, recent results from the Phase 1/2a trial revealed no significant improvement in patients, suggesting that bulk removal of toxic cholesterol is not sufficient to correct disease in the clinical setting. This is not unlike many animal disease models in which therapeutics fail to translate efficiently to the clinic resulting in failure of most drug candidates in early or late clinical trials. A deeper understanding of disease responsive features could provide considerable benefit to transitioning candidate pharmaceuticals to the clinic.

Many post-translational modifications have been the subject of therapeutic development to selectively modulate the expression and stability of disease-associated proteins. Among them acetylation has gained significant interest with the advent of small molecule regulators of histone acyl transferases (HAT) (42-44) and histone deacetylases (HDAC) (45,46), which are responsible for the reversible post-translational acetylation and deacetylation reactions of histones and non-histone proteins. There are 18 HDACs organized into four classes based on their mechanism of action including the Zn^2+^-dependent Class I, II and IV HDACs which are the target of the HDAC inhibitors (HDACi) Vorinostat (SAHA) (47), Panobinostat (LBH589) (48) and Romidepsin (FK228) (49) and the NAD+-dependent class III, sirtuins (50). While they have been extensively characterized for their role in chromatin remodelling, thereby regulating the expression of select genes, they have also been shown to regulate the post-translational acetylation state of numerous non-histone proteins, including the cytosolic chaperone Hsp90 (51,52), a key component in the regulation of the heat shock response (HSR) pathway (53-56). We have proposed that HDACis are proteostasis regulators (PR) that globally modulate cellular protein stability and function (57).

There are currently 4 HDACi’s approved by the FDA for clinical use. These include high affinity HDACis such as Panobinostat (LBH589) for the treatment of both cutaneous T-cell lymphoma (CTCL) and multiple myeloma, Belinostat (PXD101) for the treatment of Peripheral T-Cell lymphoma (PTCL) and Vorinostat (SAHA) and Romidepsin (FK228) for the treatment of CTCL (58). We have recently shown that cholesterol homeostasis in cells expressing different NPC1 variants are differentially impacted by SAHA and Panobinostat (18). These results support previous observations that the most common NPC1 variant, I1061T, is sensitive to high affinity HDACi in yeast, fibroblasts and mice (59-61), although recent results challenge this observation in the context of HPβCD treatment in a I1061T mouse model (62). HDACis have also been shown to provide substantial benefit in models of type II diabetes (46,63,64), cancer (65-67), rheumatoid arthritis (68), chronic lung diseases, including alpha-1-antitrypsin deficiency (AATD)(69-72), chronic obstructive pulmonary disease (COPD) (72-74), asthma associated airway inflammation (75-79) and cystic fibrosis (CF) (80-82).

In addition to these high affinity HDACis, the short-chained fatty acid, valproic acid (VPA) (**Figure 1B**), is a low affinity broad spectrum HDACi (class I, II & IV), which readily crosses the blood-brain-barrier (BBB). VPA has been used clinically for decades as an anticonvulsant and mood-stabilizing drug in patients suffering from epilepsy and bipolar disorder (83-85). Unlike high affinity HDACi, VPA readily crosses the BBB and exhibits a half-life of 9-18 h (86) with an improved pharmacokinetic profile compared to other HDACi. We reasoned that the characteristics that make it an effective anticonvulsant could be beneficial in the treatment of chronic neurological disorders such as NPC1, particularly given recent clinical studies of HPβCD treated patients that showed slowing of neuropsychological outcomes over a 36 month time-frame (87). While VPA is prescribed to control neurological symptoms in advanced NPC1 disease, no information is available as to its impact on the basic defect associated with the onset of NPC1 disease, namely the trafficking and functional defects associated with NPC1 variants. Given the recent failure of the VTS-270 trial (https://clinicaltrials.gov/ct2/show/NCT03879655), an incentive now exists to identify new therapeutics that can work in combination with HPβCD to improve the impact of its bulk cholesterol extraction in humans, a non-specific event which we anticipate will require improvement of NPC1 variant folding, stabilization and transport to the LE/Ly compartment to more effectively restore cholesterol homeostasis and reduce neuropsychological symptoms in the HPβCD moderated environment. Thus, the opportunity presented by the FDA approved, BBB penetrant VPA, which is administered orally (88), could provide significant benefit to existing and future patients.

In the present study, we examined the impact of VPA on NPC1-I1061T trafficking and how it affects cholesterol homeostasis. Herein, we show that VPA not only corrects the trafficking defect associated with the I1061T variant but also induced a significant reduction in LE/Ly localized CH. We also observed that combining VPA with the FDA-approved anti-malarial, chloroquine (CQ) (89), a lysosomotropic compound (90), causes a synergistic correction of the trafficking and functional defects associated with I1061T-NPC1. Our current study provides mechanistic insights and highlights the potential importance of BBB penetrant HDACis in managing proteostasis-based post-translational modifications that are responsible for promoting the trafficking of NPC1 to LE/Ly.

We posit that a combination of VPA-mediated correction of NPC1 trafficking and CQ-mediated alteration of lysosomal pH, that affects rescued NPC1 variant function, may provide a more favorable LE/Ly environment that could have clinical value in improving HPβCD based therapies. Because HPβCD based therapies only impact the toxic build-up of cholesterol in cells, they fail to address the fundamental problem in disease which will require restoration of a functional NPC1 fold in the context of the genetic diversity found in the patient population(17,18). We posit that in response to the reduced cholesterol load generated by HPβCD, pharmaceuticals that boost NPC1 variant function will be necessary to restore cholesterol recycling and downstream neuropsychological activity impacting patients.

## Results

### Valproic acid reduces cholesterol accumulation in I1061T-NPC1 expressing fibroblasts

Early experiments provided evidence that the HDACi, SAHA, can improve cholesterol clearance in primary fibroblasts homozygous for the I1061T variant (91). We have recently extended these observations and shown that the FDA-approved SAHA and LBH589 can provide functional cholesterol correction to over 68 NPC1 variants (18). While these results suggest that HDACis can restore CH homeostasis in patient-derived fibroblasts and heterologous models (59-61), our screen did not include the BBB penetrant HDACi, valproic acid (VPA) (**Fig 1B**). We first analyzed the effect of VPA on CH homeostasis in patient-derived NPC1 fibroblasts homozygous for the I1061T variant (GM18453) and a compound heterozygote (P237S/I1061T) (GM3123). The level of free, unesterified CH was quantified using the CH binding dye filipin combined with an automated microscopy analysis, as previously described (92) (**Fig. 1C & D**). Patient fibroblasts that are homozygous or heterozygous for the disease-associated I1061T variant exhibit CH accumulation relative to that seen in fibroblasts derived from WT-NPC1 expressing individuals (GM05659) (**Fig. 1C & D**), consistent with the reported NPC1 patient phenotype. Treatment of patient-derived fibroblasts with 4 mM VPA for 48 h led to a global reduction in the LE/Ly CH pool, consistent with the reported beneficial effect of other HDACis on CH homeostasis in cells expressing multiple NPC1 disease variants (18,59,60).

### VPA corrects the trafficking of I1061T-NPC1 in patient-derived fibroblasts

Human NPC1 is primarily localized to the limiting membrane of the LE/Ly compartments (**Fig. 1A**). The I1061T mutation in NPC1 results in defective folding and ER retention leading to ER-associated degradation (ERAD) through conventional and MARCH6-dependent (33,93) and ER-phagy pathways (93), generating a loss-of-function phenotype, and providing an explanation for the observed loss of cholesterol processing in the LE/Ly compartment. We observed that treating cells with increasing doses of VPA led to a dose-dependent increase in I1061T-NPC1 mRNA, reaching statistical significance at a dose of 4mM (**Fig. 2A**). Despite a nearly 4-fold increase in mRNA (**Fig. 2A**), we only observed a modest 2-fold increase in NPC1 protein at 4 mM VPA (**Fig. 2B & C**) likely reflecting its instability in the ER (33).

**Figure 2.**
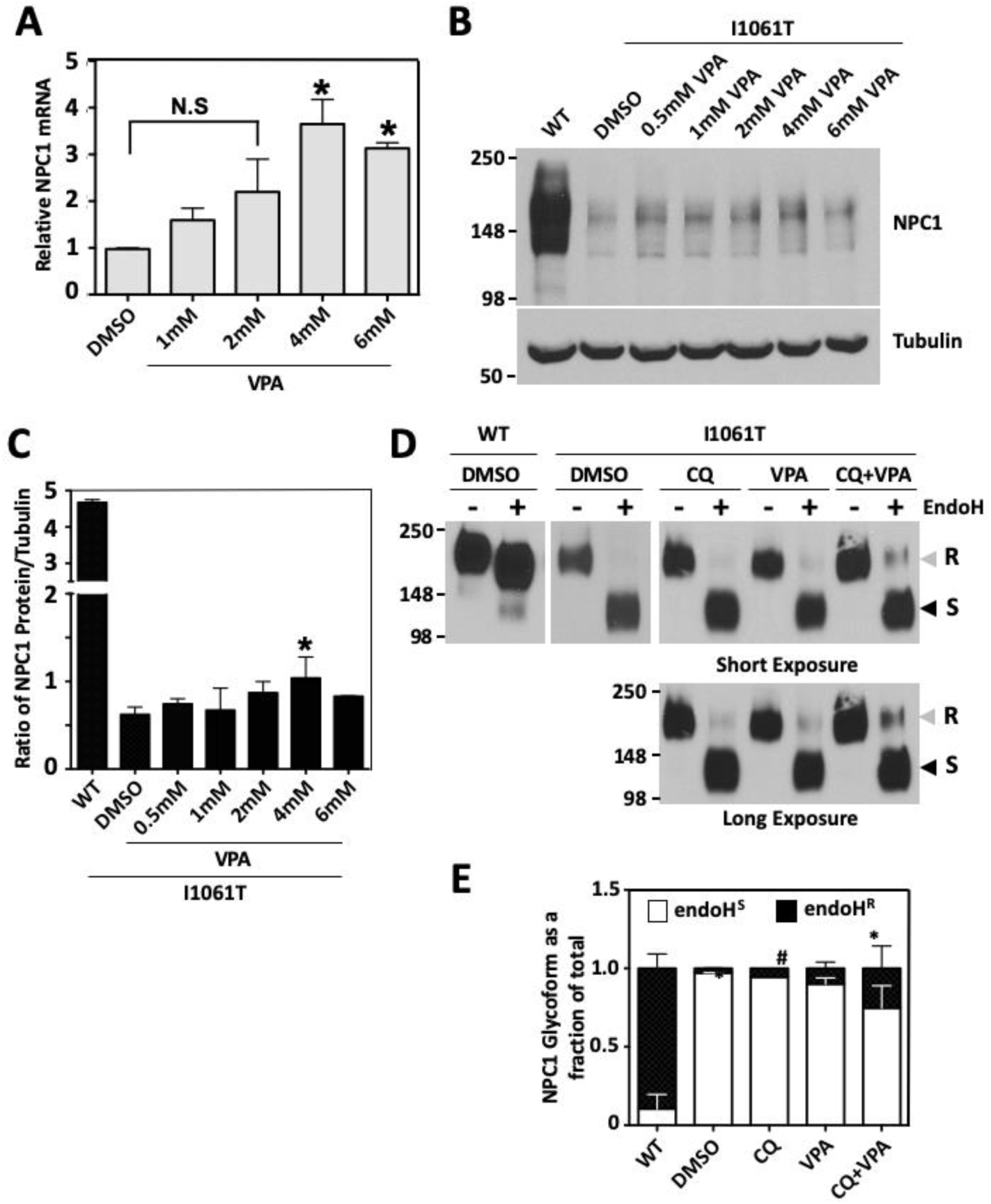
VPA enhances NPC1-I1061T expression and trafficking in human fibroblasts. **A.** Bar graph depicting the level of NPC1 mRNA in human fibroblasts homozygous for NPC1-I1061T in response to the indicated dose of VPA for 48h. The data is shown as the relative mean ± SD with the vehicle treatment normalized to 1 and asterisks indicates p < 0.05 as determined by a one-way ANOVA and Dunnett’s post-tests. **B.** Representative Western blot of NPC1 and tubulin in response to the treatment of human fibroblasts homozygous for NPC1-I1061T with the indicated dose of VPA for 48h. As a control, an equivalent total protein sample of DMSO treated WT fibroblast was included. **C.** Bar graph depicting the levels of NPC1 protein in human fibroblasts homozygous for NPC1-I1061T in response to the indicated dose of VPA for 48h. The data represents the mean ± SD of the ratio of the NPC1 to Tubulin. Asterisk indicate p < 0.05 as determined by a one-way analysis of variance and Dunnett’s post-test (n = 3). **D.** Representative Western blot of NPC1 from human fibroblasts treated with DMSO, CQ (50μM), VPA (4mM), or CQ+VPA (50μM + 4mM) for 48 h. NPC1 was immunoprecipitated and subsequently treated without (-) or with (+) Endo H prior to Western blot analysis. **E.** Bar graph depicting the amount of EndoH^R^ (white) and EndoH^R^ (black) glycoforms as a fraction of total NPC1 for WT- and I1061T-NPC1 treated as in panel **D**. Asterisk indicate p < 0.05 as determined by a one-way analysis of variance and Dunnett’s post-test using DMSO treated I1061T as the reference and hashtag indicates p < 0.05 as determined by a one-way analysis of variance and Dunnett’s post-test using VPA treated I1061T as the reference (n = 3).

To assess if the increased expression correlated with increased trafficking of the I1061T ER-restricted variant, we monitored the trafficking of NPC1 following the treatment of the I1061T variant GM18453 fibroblasts with a dose of 4mM VPA for 48h **(Fig. 2D & 2E**). Human NPC1 contains 14 N-linked glycans that are added co-translationally during translocation into the ER (94). The trafficking of NPC1 from the ER to the LE/Ly compartment can be monitored by Western blot analysis (**Figure 2D**) where the post-ER fraction of NPC1 acquires resistance (R) to endoglycosidase H (Endo H) due to N-glycan modifications in the Golgi leading to slower migration on SDS-PAGE. At steady state, nearly 80% of the wild-type (WT) NPC1 exhibits EndoH^R^ **(Fig. 2D & 2E, WT + DMSO**), whereas less then 10% of the I1061T variant exhibits EndoH^R^ (**Fig. 2D & 2E, I1061T + DMSO**). The treatment of GM18453 fibroblasts with 4mM VPA resulted in a 2-fold increase in the EndoH^R^ fraction of I1061T-NPC1 (**Fig. 2D & 2E**). Although VPA did provide a statistically significant correction of the trafficking defect associated with the I1061T variant, the amount of NPC1 reaching the LE/Ly compartment was disproportionate to the observed effect on CH clearance, where we observed a > 60% reduction (**Fig. 1C**) suggesting that a modest effect on I1061T trafficking can have a major effect on restoration of cholesterol homeostasis or that the LE/Ly-localized fraction of I1061T-NPC1 is destabilized upon reaching the LE/Ly compartment.

HDACi have been shown to induce a hyper-acidification of the lysosome compartment (95,96), which is likely to alter lysosomal function. One possibility to account for the discrepancy between the level of I1061T-NPC1 trafficking and LE/Ly CH reduction may arise because of a destabilization of VPA-corrected NPC1-I1061T variant protein in the lysosomes. To test this hypothesis, we co-treated I1061T-expressing GM18453 fibroblasts with 4mM VPA in the presence of CQ, a compound known to partially neutralize lysosomal acidification (97). While CQ treatment alone had no effect on the trafficking of NPC1 (**Fig. 2D & 2E**), the combined VPA + CQ treatment provided a synergistic 4-fold increase in I1061T-associated EndoH^R^ (**Fig. 2D & 2E**) relative to that seen with either treatment alone.

### CQ neutralizes the HDACi-mediated acidification of lysosomes in human fibroblasts

It has previously been demonstrated that NPC1 exhibits a transmembrane cation efflux pump activity capable of removing acriflavine, a cationic dye, from the lysosomal compartment contributing to lysosomal acidification (98). Thus, the absence of LE/Ly-localized NPC1 in GM18453 fibroblasts could contribute to the alkalization of the organelle. Because CQ can synergistically improve the VPA-corrected post-ER fraction of I1061T-NPC1 in patient derived fibroblasts, we hypothesized that the effect of CQ is to neutralize the joint VPA and I1061T-mediated hyper-acidification of the lysosomal compartment and therefore lead to improved stability of I1061T-NPC1.

To test this hypothesis, we monitored the acidification of lysosomes in both WT and I1061T patient-derived fibroblasts in response to 4mM VPA (**Fig. 3**). Lysosomal pH can be monitored in vivo using acridine orange, a metachromatic, weak base dye that stains lysosomes and nucleic acids (99). In the acidic conditions typically observed in the lysosome, the acridine dye becomes protonated and accumulates within the organelle where it forms a precipitate that emits a red-shifted fluorescence. In WT fibroblasts (GM05659), we observed this red-shifted fluorescence indicative of the acidic pH of the lysosome (**Fig. 3A, panels a-c)**. Conversely, I1061T-expressing fibroblast (GM18453) exhibit a green-shifted fluorescence (**Fig. 3A, panels d-f**), indicative of alkalization of the lysosome. The treatment of the I1061T-expressing fibroblasts with VPA led to the re-acidification of the lysosome (**Fig. 3B, panels d-f**), exemplified by an increase in red fluorescence, and a concomitant decrease in green fluorescence. The effect of VPA on the acidification of the lysosome in I1061T-expressing fibroblasts is in agreement with the effect of HDACi on lysosomal pH (95,96). We also observed that the treatment of both WT-(GM05659) and I1061T-expressing (GM18453) fibroblasts with CQ, lead to a green shift in the acridine orange fluorescence, indicative of alkalization of the lysosomal compartment (**Fig. 3C**), in agreement with the reported effect of this small molecule (97).

**Figure 3.**
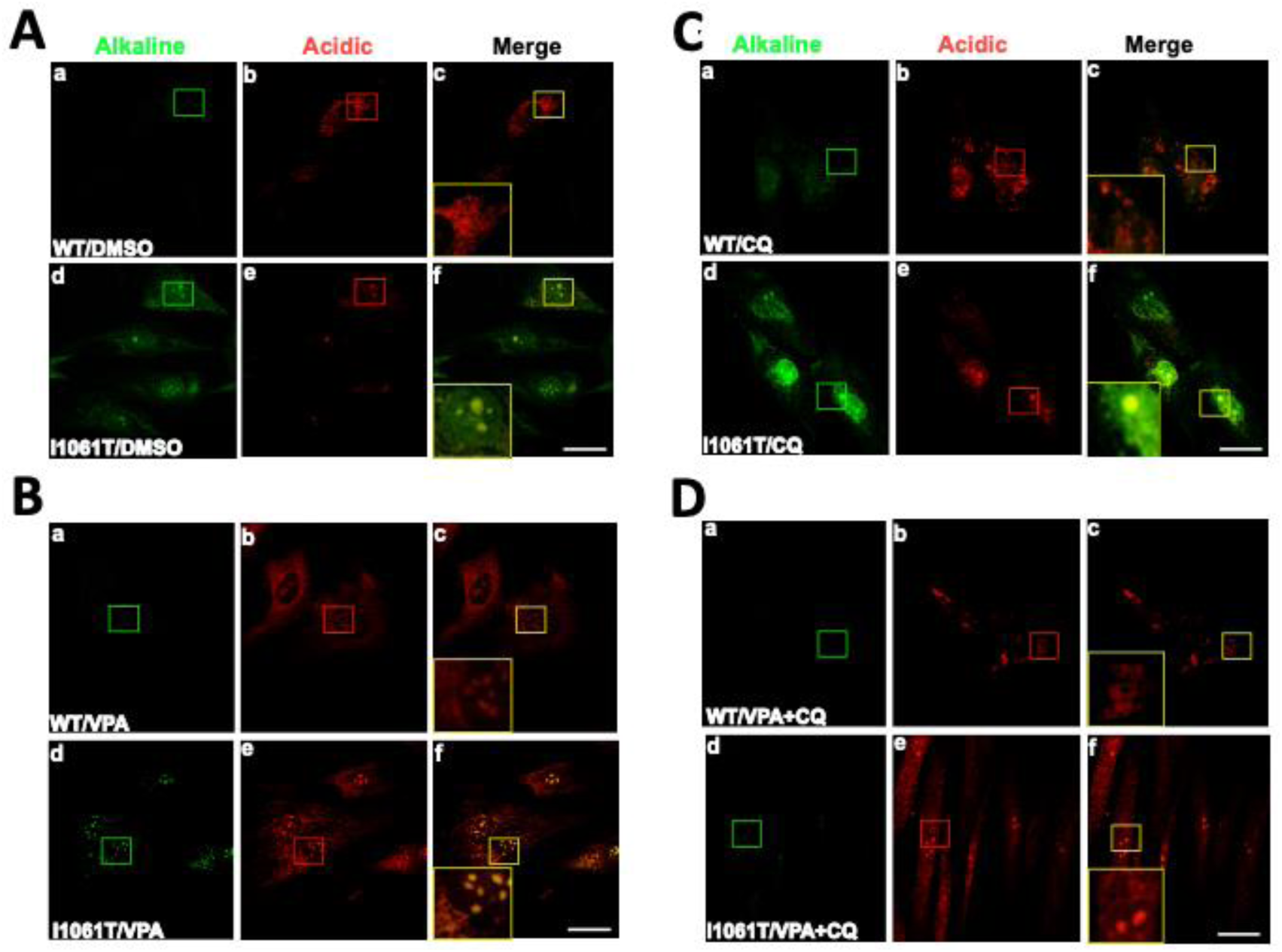
Chloroquine neutralizes the VPA-mediated acidification of the lysosomal. Representative epifluorescent images of human fibroblasts homozygous for WT-(GM05659) or I1061T-NPC1 (GM18453) stained with acridine orange (Ex/Em = 460/620) following the treatment with DMSO (**A**), 4mM VPA (**B**), 50μM CQ (**C**) or 4mM VPA + 50μM CQ (**D**) for 48 h. Scale bar represents 10 μm)

When we monitored the impact of combining the treatments of VPA and CQ on the pH of the lysosomal compartment we observed a decrease in the alkalization of the lysosome relative to that seen in I1061T-expressing cells treated with CQ alone, indicating that the expected CQ-mediated alkaline pH shift was countered by the acidifying effect of VPA **(Fig. 3D, panels d-f).** These data confirm our hypothesis above, that while VPA promoted ER export of I1061T, its effect on lysosomal pH resulted in an unstable lysosomal pool of NPC1. CQ mitigated the VPA-mediated acidification of the lysosomal compartment resulting in the accumulation of a stable I1061T-associated EndoH^R^ fraction.

### Combining VPA and CQ synergistically corrects the trafficking of I1061T-NPC1 in HeLa cells

To explore the combinatorial effect of VPA and CQ on NPC1 rescue, we established a HeLa cell culture model that replicates the NPC1 phenotype (**Fig. S1**). Because NPC1 is ubiquitously expressed, we first generated NPC1 null HeLa cells using a lentiviral shRNA targeting the 3’-UTR of NPC1 (**Fig. S1A-C**). WT- and I1061T-HeLa cells were generated by transducing the NPC1 null HeLa cells with lentivirus delivering the appropriate NPC1 variant cDNA (**Fig. S1C**). An analysis of the trafficking revealed a fractional distribution of EndoH^R^ glycoforms that is consistent with that seen in patient derived fibroblasts for each of these NPC1 variants (**Fig. S1C & S1D**). We also used immunofluorescence to confirm the subcellular localization of these NPC1 variants in HeLa-shNPC1 cells (**Fig. S2A-C**). Here we noted that WT-NPC1 had a punctate staining pattern that co-localizes extensively with the lysosomal marker LAMP1 (**Fig. S2A, panels a-c**) and exhibited little to no co-localization with the ER markers TMEM97 (100) (**Fig. S2B, panels a-c**) and KDEL (101) (**Fig. S2C, panels a-c**). Conversely, I1061T-NPC1 exhibited a reticular staining pattern that did not co-localize with LAMP1 (**Fig. S2A, panels e-g**) but rather exhibited significant overlap with TMEM97 (**Fig. S2B, panels d-f**) and KDEL (**Fig. S2C, panels d-f**). Additionally, we observed a diffuse filipin staining pattern in WT-NPC1 expressing HeLa-shNPC1 cells (**Fig. S2A, panel d**), whereas filipin positive perinuclear puncta where observed in I1061T-expressing cells (**Fig. S2A, panel h**). These data are consistent with the respective cholesterol homeostasis observed in WT- and I1061T-NPC1 expressing primary fibroblasts (**Fig. 1C**).

With the establishment of WT- and I1061T-HeLa cells we determined if these cell lines recapitulated the VPA/CQ-mediated correction of the I1061T variant seen in patient derived fibroblasts (**Fig. 1 & 2**). The treatment of HeLa-I1061T cells with VPA resulted in the appearance of an EndoH^R^ pool of NPC1-I1061T reaching 10% of total at a dose of 4mM VPA (**Fig. 4A & B**), a level of correction consistent with that seen in the I1061T homozygote patient-derived fibroblasts (GM05659) (**Fig. 2D & E)**. We also observed a dose-dependent increase in the mRNA of I1061T-NPC1, which saturated between 2-6mM (**Fig. 4C**), similar to what we observed for the impact of VPA on GM05659 fibroblasts (**Fig. 2A**). To ascertain if the lysosomotropic CQ would synergize with VPA to further increase the trafficking of the stably expressed I1061T-NPC1 in HeLa-shNPC1 cells, we combined these compounds and found that while VPA increased the amount of EndoH^R^ I1061T-NPC1 to 10% of total, the addition of CQ improved this correction to 50% of total (**Fig. 4D & E**), a result consistent with the synergistic correction of this disease variant in patient derived fibroblasts (**Fig. 2D & E**).

**Figure 4.**
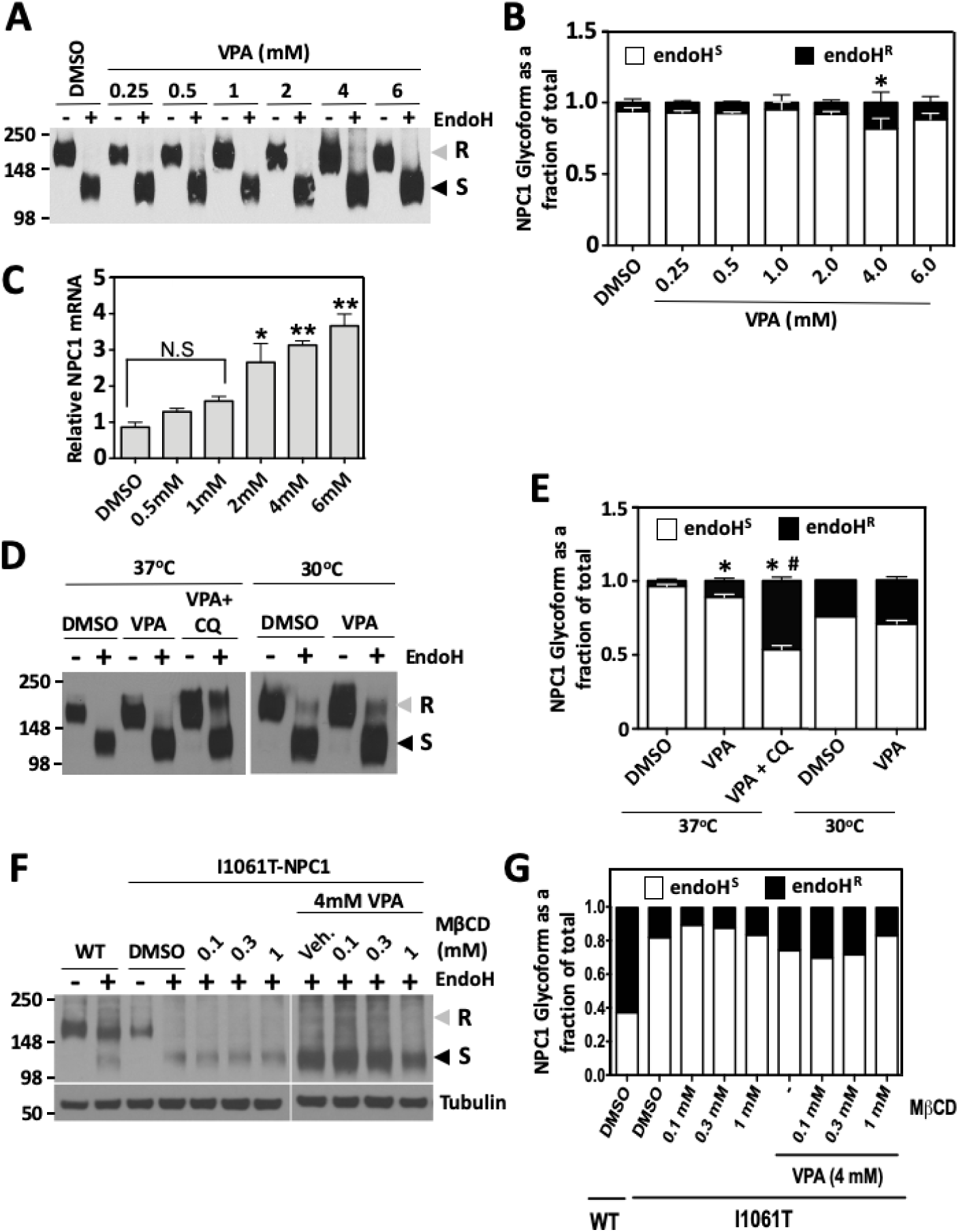
VPA corrects I1061T-NPC1 trafficking in HeLa-shNPC1 cells. **A.** Representative Western blot of I1061T-expressing HeLa-shNPC1 cell lysates treated with the indicated concentrations of VPA for 24 h. NPC1 was immunoprecipitated and treated without (-) or with (+) Endo H prior to SDS-PAGE. **B.** Bar graph depicting the amount of EndoH^S^ (white) and EndoH^R^ (black) glycoforms as a fraction of total NPC1 from I1061T-expressing HeLa-shNPC1 cell lysates treated with the indicated concentrations of VPA for 24 h. The data are presented as the normalized mean ± SD and the asterisk indicates p < 0.05 using a two-tailed T-test with DMSO treatment as a reference (n = 3). **C.** Bar graph depicting the NPC1 mRNA levels in I1061T-expressing HeLa-shNPC1 cell lysates treated with the indicated concentrations of VPA for 24 h. The data is shown as the relative mean ± SD with the vehicle treatment normalized to 1 and asterisks indicates p < 0.05 and double asterisk indicating p < 0.01 and N.S indicating not significant as determined by a one-way ANOVA and Dunnett’s post-tests (n = 3). **D.** Representative Western blot of I1061T-expressing HeLa-shNPC1 cell lysates treated with DMSO, VPA (4mM), or CQ+VPA (50μM + 4mM) for 24 h or incubated at 30°C in the presence of DMSO or 4mM VPA for 24h. NPC1 was immunoprecipitated and treated without (-) or with (+) Endo H prior to SDS-PAGE. **E.** Bar graph depicting the amount of EndoH^S^ (white) and EndoH^R^ (black) glycoforms as a fraction of total NPC1 from I1061T-expressing HeLa-shNPC1 cell lysates treated with the indicated treatment for 24 h as in panel **D**. The data are presented as the normalized mean ± SD and the asterisk indicates p < 0.05 using a two-tailed T-test with DMSO treatment as a reference and the hashtag represents p < 0.05 using a two-tailed T-test with VPA treatment as a reference (n = 3). **F.** Western blot of WT- or I1061T-expressing HeLa-shNPC1 cell lysates treated with DMSO, 4mM VPA, or the indicated concentration of methyl-β-cyclodextrin (MβCD) for 24 h. NPC1 was immunoprecipitated and treated without (-) or with (+) Endo H prior to SDS-PAGE. **G.** Bar graph depicting the amount of EndoH^S^ (white) and EndoH^R^ (black) glycoforms as a fraction of total NPC1 from WT- and I1061T-expressing HeLa-shNPC1 cell lysates treated as in panel **F**.

The misfolding and stability of I1061T-NPC1 arises due to the inability of the variant polypeptide chain to properly engage key component(s) of the prevailing proteostasis environment. The inability to achieve a WT-like fold results in a loss of protein stability leading to its degradation by conventional and MARCH6-dependent (33,93) ERAD and ER-phagy pathways (93). Previous studies have shown that incubation of cells expressing meta-stable proteins, such as the F508del variant of CFTR (102-104) and NPC1 I1061T (33) at low temperature (27°-30°C) can improve protein folding and stability leading to a partial correction of the disease phenotype. The exposure of I1061T-HeLa cells to 30°C for 24 h led to a partial correction of the trafficking defect associated with this disease variant, with the EndoH^R^ fraction reaching 25% of total (**Fig. 4D, E**). Combining the low temperature correction with VPA treatment did not provide significant improvement in the trafficking of I1061T relative to that seen at 30°C alone (**Fig. 4D & E**). This observation suggests that the effect of VPA treatment on the I1061T-NPC1 protein is similar to the effect of low temperature, a condition that improves the folding and stability of the variant protein to achieve egress from the ER for CQ mediated stabilization in the LE/Ly compartment.

Recent data has highlighted the benefits of HPβCD, a cholesterol binding compound that extracts recycling cholesterol from endomembrane systems at the cell surface. HPβCD has been shown to reduce both unesterified cholesterol and glycolipids, thereby prolonging survival and delaying the onset of the neurodegenerative phenotype in preclinical mouse models (39,41,105). Since, cyclodextrins have been shown to correct the lipid accumulation in NPC1 null mice, its action is independent of NPC1 as would be expected of a bulk phase lipid-binding compound. In order to ascertain the effect of cyclodextrin on the trafficking of I1061T-NPC1, we monitored the effect of different concentrations of methyl-β-cyclodextrin (MβCD) alone and in combination with VPA on the extent of I1061T-EndoH^R^ in I1061T-expressing HeLa-shNPC1 cells. MβCD alone had no effect on the trafficking of I1061T-NPC1 (**Fig. 4F & 4G**), however, we did observe a slight improvement in the VPA-mediated correction of I1061T at lower doses of HPβCD (0.1mM) (**Fig. 4F & 4G**). Interestingly increasing the concentration of MβCD appeared to abrogate this effect, where at 0.3mM MβCD we see a return to VPA-mediated trafficking alone and at 1mM MβCD we see 50% decrease in the extent of EndoH^R^ level seen with VPA alone (**Fig. 4F & 4G**). These data suggest these compounds might be acting on competing pathways to exert their effect on cells, which, when pushed too hard, result in inhibitory effects on the trafficking of I1061T-NPC1 rather than a synergistic corrective effect. These results reflect the expected dynamic interplay between the composition of the lipid bilayer of endomembrane compartments that form a gradient of cholesterol with low cholesterol in the ER and high cholesterol (50%) in the plasma membrane influencing NPC1 trafficking and function in unknown ways.

### Valpromide does not correct the trafficking of I1061T-NPC1

Valpromide (VPM) is a VPA analog in which the carboxyl group has been modified to an amide group (**Fig. S3A**) thereby removing its deacetylase inhibitory activity (106,107). Despite the loss of this enzymatic activity, VPM still maintains its anticonvulsant properties (108). To assess the contribution of the deacetylase activity of VPA in the correction of the trafficking and functional defect of NPC1-I1061T, we tested the ability of VPM to mediate correction of the I1061T variant in HeLa-shNPC1 cells. VPM failed to increase NPC1 mRNA (**Fig. S3B**) or increase EndoH^R^ glycoform of the I1061T variant either alone or in combination with CQ (**Fig. S3C & S3D**), suggesting that the deacetylase activity of VPA is critical for its corrective properties. Interestingly, we observed a dose-dependent competitive inhibition by VPM on the VPA/CQ-mediated correction of I1061T-NPC1 (**Fig. S3E & S3F**). These data suggest that while VPM has lost its deacetylase activity, it maintains its capacity to bind to key regulatory components involved in the VPA-mediated correction of I1061T-NPC1, thereby inhibiting the corrector activity of VPA.

### The combination of VPA and CQ stabilizes NPC1-I1061T

Given the improvement in trafficking seen with the treatment of VPA in the absence and presence of CQ, we performed a pulse chase analysis of WT- and I1061T-NPC1 to determine if any of these treatments increased the stability of the I1061T variant in the ER and post-ER compartments. Under control conditions ∼20% of WT-NPC1 is degraded over the 12h chase period (**Fig. 5A**) revealing that WT NPC1 is both stable and efficiently trafficked from the ER, an observation supported by the fact that at T=0 we detect an EndoH^R^ fraction representing 35% of total which more than doubles at T=12h (**Fig. 5A**). An analysis of the DMSO treated I1061T variant reveals a decrease in protein stability relative to that seen with WT-NPC1 and failure to accumulate the EndoH^R^ glycoform (**Fig. 5B**). We noted that CQ had no effect on the stability or maturation of I1061T-NPC1 in the ER (**Fig. 5C**), consistent with our observations above that it did not correct the trafficking defect linked with the I1061T variant. While VPA did show an increased accumulation of the EndoH^R^ glycoform at all time points (**Fig. 5D**), consistent with its ability to correct the I1061T-associated trafficking defect (**Fig. 2D, 4A & 4D**), it did not improve the stability of the EndoH^S^ glycoform, showing an ∼60% reduction in this ER fraction (**Fig. 5D**), a level similar to that seen with DMSO treated I1061T-NPC1. Despite the inability of either VPA or CQ to independently provide a significant increase in the stability of I1061T-NPC1, the combination of these 2 small molecules caused a significant improvement in the stability of this disease-associated variant, resulting in a WT-like 20% reduction in total I1061T NPC1 relative to the level seen at T = 0 (**Fig. 5E**). Notably, the entirety of the remaining I1061T-NPC1 at T = 12h is recovered in the EndoH^R^ glycoform (**Fig. 5E**), indicating that the trafficking of the I1061T variant in the VPA+CQ modified environment is very efficient. Additionally, these data suggest that the EndoH^S^ glycoform seen at steady state after the treatment with VPA and CQ represent either a *de novo* synthesized pool that has yet to traffic out of the ER, or an accumulation of a non-rescuable I1061T-NPC1 pool that was synthesized prior to the addition of the correctors.

**Figure 5.**
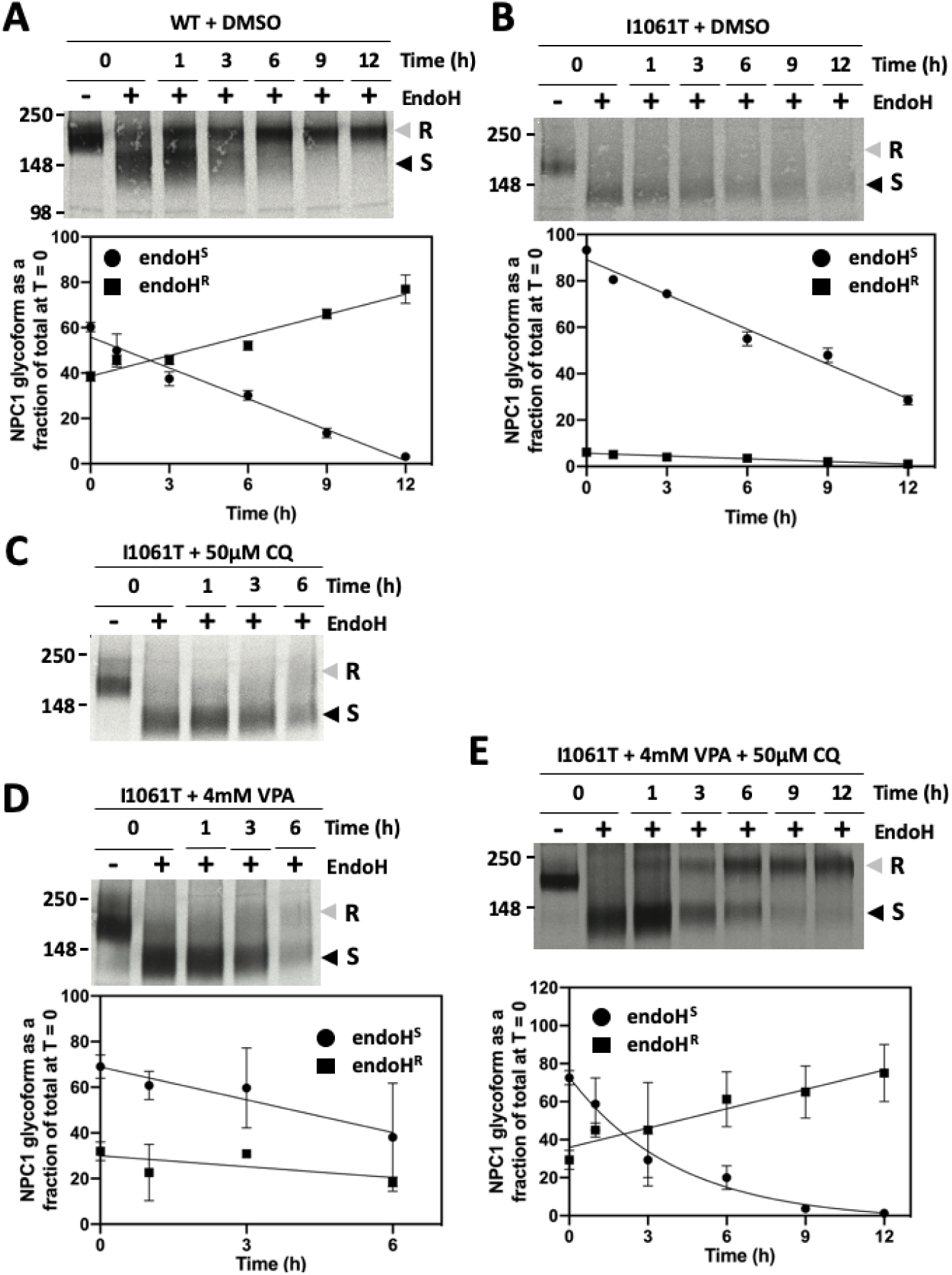
Chloroquine increases the stability of VPA-corrected I1061T-NPC1. **A. (Top)** Autoradiograph of pulse labeled WT-NPC1 stably expressed in HeLa-shNPC1 cells treated with DMSO for 24h and chased for the indicated time. **(Bottom)** Scatter plot depicting the fractional distribution of EndoH^S^ (circle) and EndoH^R^ (square), normalized to the amount of total NPC1 at T = 0, over the chase period. **B. (Top)** Autoradiograph of pulse labeled I1061T-NPC1 stably expressed in HeLa-shNPC1 cells treated with DMSO for 24h and chased for the indicated time. **(Bottom)** Scatter plot depicting the fractional distribution of EndoH^S^ (circle) and EndoH^R^ (square), normalized to the amount of total NPC1 at T = 0, over the chase period. **C.** Autoradiograph of pulse labeled I1061T-NPC1 stably expressed in HeLa-shNPC1 cells treated with 50μM CQ for 24h and chased for the indicated time. **D. (Top)** Autoradiograph of pulse labeled I1061T-NPC1 stably expressed in HeLa-shNPC1 cells treated with 4mM VPA for 24h and chased for the indicated time. **(Bottom)** Scatter plot depicting the fractional distribution of EndoH^S^ (circle) and EndoH^R^ (square), normalized to the amount of total NPC1 at T = 0, over the chase period. **E. (Top)** Autoradiograph of pulse labeled I1061T-NPC1 stably expressed in HeLa-shNPC1 cells treated with 4mM VPA + 50μM CQ for 24h and chased for the indicated time. **(Bottom)** Scatter plot depicting the fractional distribution of EndoH^S^ (circle) and EndoH^R^ (square), normalized to the amount of total NPC1 at T = 0, over the chase period.

### VPA causes a hyperacetylation of I1061T-NPC1

While HDACi are able to post-translationally modify both histone and non-histone proteins to change the composition and/or functionality of the proteostasis environment, it is also possible that part of the VPA-mediated correction of the I1061T trafficking defect is associated with a direct hyperacetylation of NPC1. To address this possibility, we immunoprecipitated all acetylated proteins with an acetylated-lysine (AcK) antibody and monitored the extent of NPC1 recovered following treatment with VPA or vehicle (**Fig. 6A**). We observed that I1061T-NPC1 is only weakly acetylated under control conditions (**Fig. 6A**), as only a slight increase in NPC1 was observed relative to the background signal seen in shNPC1-HeLa cells. Conversely, the addition of VPA to these I1061T-HeLa cells resulted in an increase in NPC1 acetylation (**Fig. 6A**).

**Figure 6.**
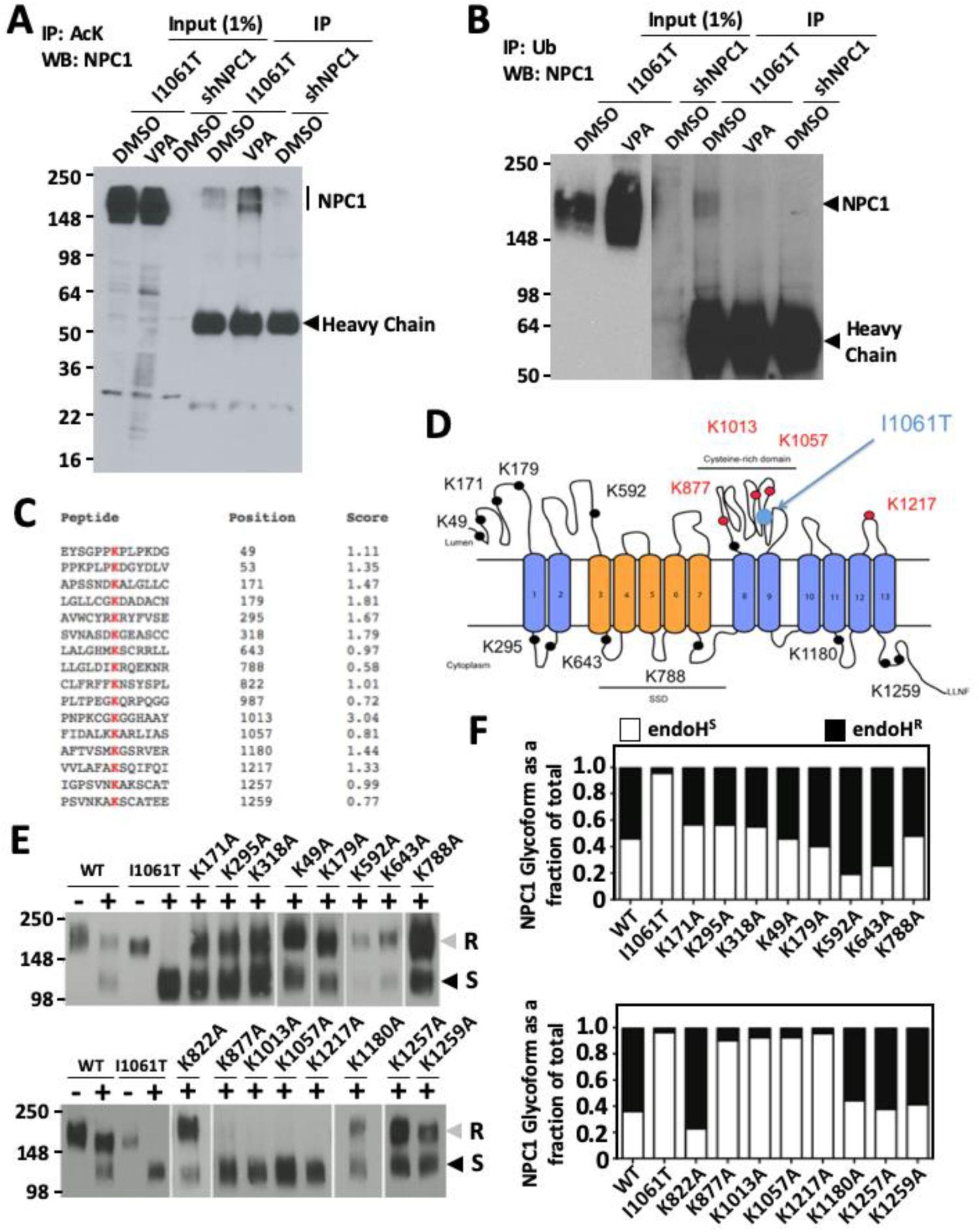
VPA mediates the hyperacetylation of I1061T-NPC1. **A.** Western blot analysis of NPC1 from immunoprecipitated (IP) acetylated lysines and associated input samples from HeLa-shNPC1 and I1061T-expressing HeLa-shNPC1 cell lysates treated with DMSO or 4mM VPA for 24h. **B.** Western blot analysis of NPC1 from immunoprecipitated (IP) ubiquitinated proteins and associated input samples from HeLa-shNPC1 and I1061T-expressing HeLa-shNPC1 cell lysates treated with DMSO or 4mM VPA for 24h. **C.** Table of predicted acetylated lysines in NPC1. Shown are the peptide sequence, the position of the predicted acetylated lysine and the associated prediction score. **D.** Cartoon of the NPC1 structure and associated domain topology showing the location of the predicted acetylated lysine shown in panel **C. E.** Western blot analysis of WT-NPC1 carrying the indicated lysine (K) mutations transiently transfected into HeLa-shNPC1 cells. Included is I1061T-NPC1 as a negative trafficking control. NPC1 was immunoprecipitated and treated without (-) or with (+) Endo H prior to SDS-PAGE. **F.** Bar graph depicting the amount of EndoH^S^ (white) and EndoH^R^ (black) glycoforms as a fraction of total NPC1 for the indicated variant transiently expressed in HeLa-shNPC1 cells. The data are presented as normalized fractions of total NPC1 in each sample.

While the acetylation of lysine residues can alter the interaction of a given protein with its normal cohort of interacting proteins (109,110), it can also block the post-translational ubiquitination of these lysine residues and thereby prevent their degradation. To address if the VPA-mediated hyperacetylation of NPC1 altered its ubiquitination, we immunoprecipitated ubiquitinated (Ub) proteins and monitored the recovery of I1061T-NPC1. Here we observed a decrease in the amount of Ub-NPC1 in lysates from VPA-treated I1061T-HeLa cells (**Fig. 6B**), indicating that part of the corrective properties of VPA are associated with the alteration in the post-translational modifications of NPC1, which promotes increased association with trafficking components and decreased association with degradative components.

To address the location of these post-translational acetylation sites, we used the Prediction of Acetylation on Internal Lysine (PAIL) algorithm to predict which residues might be targeted by acetyl transferases. The algorithm predicted 16 lysine residues would be possible acetylation targets in NPC1 (**Fig. 6C & D**). In order to assess the contribution of these lysines residues to the trafficking of NPC1, we proceeded to mutagenize these sites to alanines in WT-NPC1 and assess the trafficking ability of these variants. While we observed that most K/A mutations had no impact on the trafficking of WT-NPC1 (**Fig. 6E & 6F**), we observed that 4 lysines mutants, namely K877A, K1013A, K1057A and K1217A exhibited a complete inability to traffic out of the ER (**Fig. 6E & 6F**), as exemplified by their lack of EndoH^R^. All 4 of these lysine residues are localized on NPC1 domains that face the luminal side of cellular organelles. Interestingly, 3 of the 4 are localized in the cysteine-rich domain (CRD) of NPC1 (**Fig. 6D**), which is the site of the most prevalent disease-causing mutations, I1061T. These data suggest that the I1061T mutations might alter the folding of NPC1 allowing for access of these lysine residues to a as yet to be characterized luminal deacetylase, which is sensitive to VPA, that contributes to the trafficking defect seen with this disease-associated variant-an effect that is inhibited by VPA.

### Knockdown of HDAC7 partially restores the trafficking of NPC1-I1061T mutant in HeLa-shNPC1 cells

There have been 18 HDACs identified in the human genome belonging to two distinct families with different catalytic mechanisms, namely the Zn^2+^-dependent histone deacetylases (HDAC1-11), which are sensitive to VPA, and NAD^+^ -dependent sirtuins (SIRT1-7) (111). HDACs play regulatory roles in many biological processes, and recently emerged as promising therapeutic targets to correct the defects associated with several protein misfolding diseases (112). We have previously shown that the HDACi-mediated correction of the trafficking defect associated with the F508del variant of CFTR and the Z-variant (E342L) of alpha-1 antitrypsin (AAT), which is the primary disease-linked variant leading to AAT deficiency, occurs through the SAHA-mediated silencing of HDAC7 (71,81).

In order to determine if a similar mechanism of action is occurring for the VPA-mediated correction of I1061T-NPC1, we assessed the impact of this HDACi on the expression of HDAC7 in HeLa-shNPC1 cells stably expressing the I1061T variant. Here, we observed a dose dependent reduction in HDAC7 protein levels in response to increasing doses of VPA, with a complete loss observed at 4 mM (**Fig. 7A & B**), a dose which provides the maximal correction of the trafficking and functional defects associated with I1061T-NPC1 in HeLa and primary fibroblasts. To determine if the VPA-mediated silencing of HDAC7 is responsible for the observed correction of the I1061T variant, we used siRNA to assess the impact of silencing all class I, II and IV HDACs (HDAC 1-11) on the trafficking of I1061T-NPC1 in HeLa-shNPC1 cells. In agreement with our previous observations, we observed that the silencing of HDAC7 did significantly improve the trafficking of I1061T-NPC1, reaching a level of EndoH^R^ approaching 50% of total NPC1 (**Fig. 7C & 7D**), an effect that was further improved by the addition of CQ (**Fig. 7C & D**). We also observed that the silencing of HDAC5 and HDAC8 also provided a significant improvement of the trafficking of the I1061T variant (**Fig. 7C & 7D**), but only the siHDAC5-mediated correction was responsive to CQ (**Fig. 7C & 7D**). Despite the latter observations, the silencing of HDAC7 does appear to provide the most robust correction of the trafficking defect of I1061T-NPC1.

**Figure 7.**
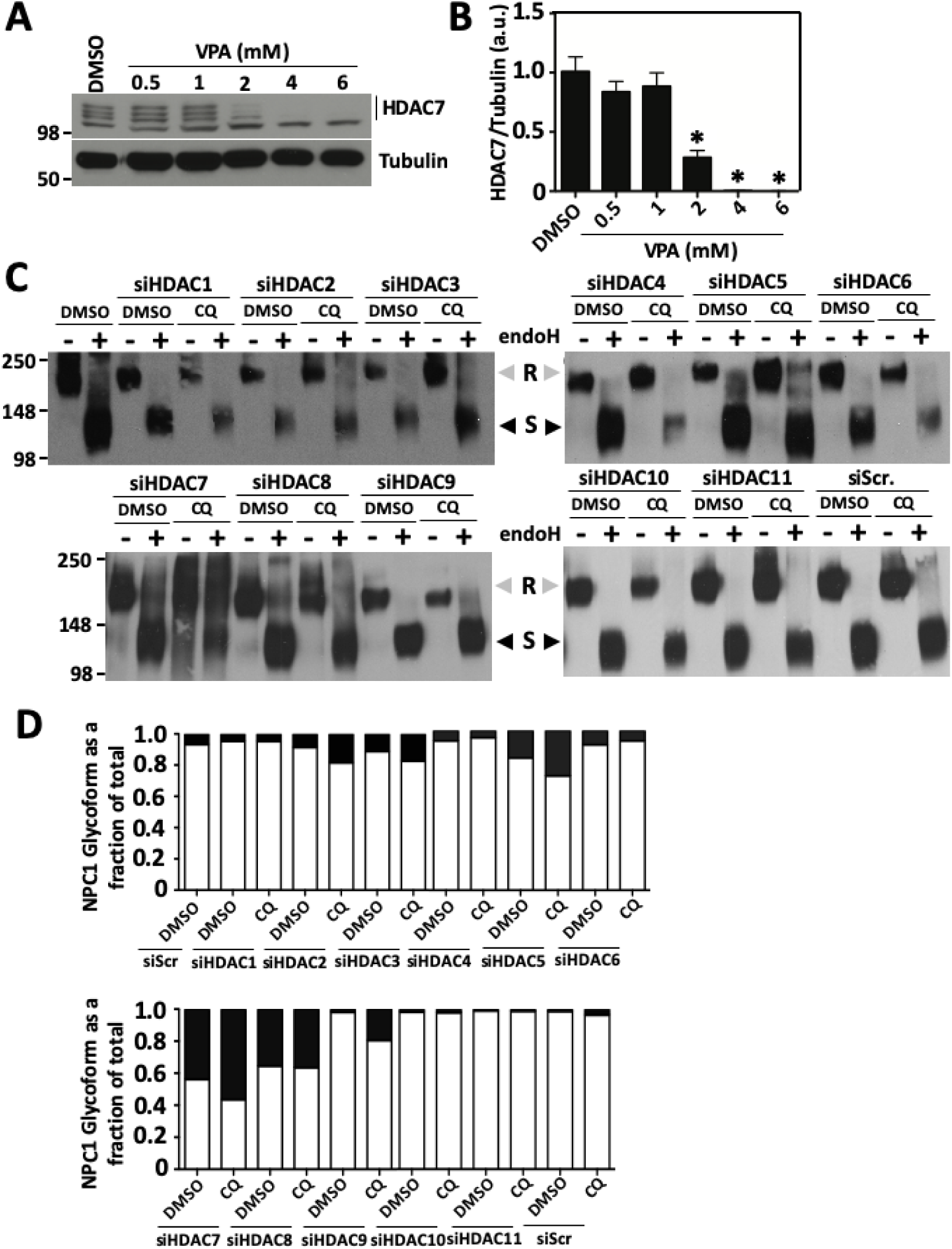
The silencing of HDAC7 corrects the trafficking of I1061T-NPC1. **A.** Representative Western blot analysis of HDAC7 and Tubulin from I1061T-expressing HeLa-shNPC1 cell lysates treated with the indicated concentrations of VPA for 24 h. **B.** Bar graph depicting the normalized ratio of HDAC7 to Tubulin in I1061T-expressing HeLa-shNPC1 cells treated with the indicated concentrations of VPA for 24 h. The data are shown as the mean ± SD normalized to 1 for the DMSO treatment condition. Asterisk represents p < 0.05 as determined by a two-tailed t-test using DMSO as the reference (n = 3). **C.** Western blot analysis of NPC1 from I1061T-expressing HeLa-shNPC1 cell lysates treated with the indicated siRNA in the absence (DMSO) or presence of 50 μM CQ. NPC1 was immunoprecipitated and treated without (-) or with (+) Endo H prior to SDS-PAGE. **D.** Bar graph depicting the level of EndoH^S^ (white) and EndoH^R^ (black) glycoforms as a fraction of total NPC1 in I1061T-expressing HeLa-shNPC1 cells treated with the indicated siRNA in the absence (DMSO) or presence of 50 μM CQ.

To confirm the that the silencing of HDAC7 corrects the trafficking defect of I1061T-NPC1, we performed immunofluorescence analysis of I1061T-NPC1 in HeLa-shNPC1 cells treated with siHDAC7. Relative to that seen in cells treated with a control siRNA (siScr) (**Fig. 8A – panels a-c**), we observed a significant increase in the co-localization of I1061T-NPC1 with the lysosomal marker, LAMP1 (**Fig. 8A – panels e-g**). The silencing of HDAC7 in I1061T-expressing HeLa-shNPC1 cells also provided a significant abrogation of the LE/Ly CH accumulation (**Fig. 8A – panel h**) compared to that seen in siScr treated cells (**Fig. 8A – panel d**). Additionally, we observed that the silencing of HDAC7 caused a >4-fold increase in the expression level of I1061T-NPC1 (**Fig. 8B & C**), suggesting that the absence of this class II HDAC improves the expression of NPC1, promotes its stability, inhibits its degradation or a combination of these effects, resulting in an improved LE/Ly localization of a functional pool of I1061T-NPC1. An examination of the impact of combining siHDAC7, with either VPA or CQ revealed that both compounds further improved the trafficking of the I1061T variant relative to that seen with siHDAC7 treatment alone (**Fig. 8D & E**). These results suggest that the VPA-mediated silencing of HDAC7 is a significant contributing factor in the mechanism of action for the VPA-mediated correction of the I1061T-NPC1 at the biochemical and molecular level.

**Figure 8.**
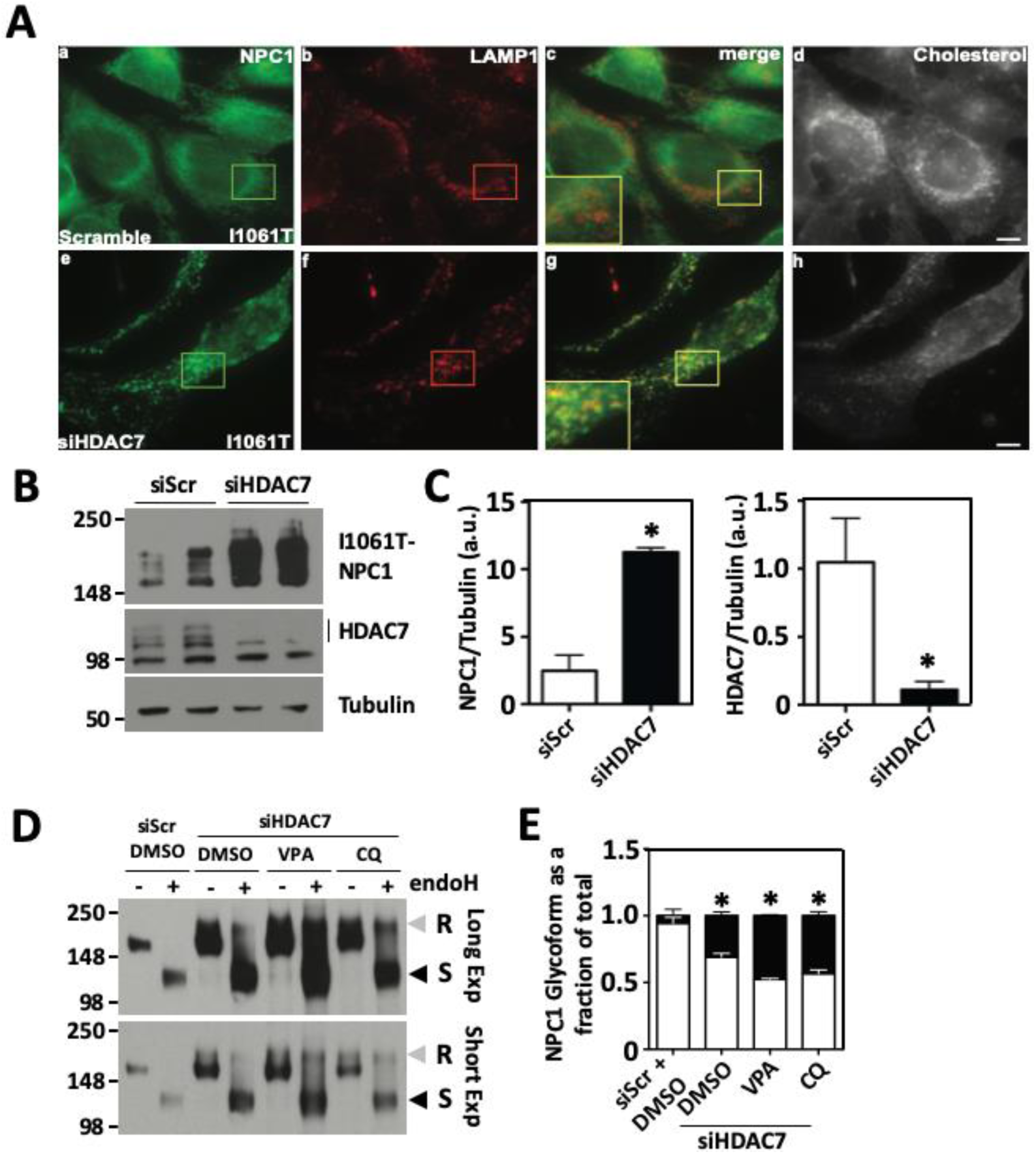
The silencing of HDAC7 restores LE/LY-localized I1061T-NPC1. **A.** Representative immunofluorescent images of I1061T-expressing HeLa-shNPC1 cells treated with control siRNA (siScr) (panels a-d) or siHDAC7 (panels e-h). Shown are NPC1 (green) (panels a & e), LAMP1 (red) (panels b & f) and filipin (greyscale) (panels d & h) staining. **B.** Representative Western blot of NPC1, HDAC7 and Tubulin from I1061T-expressing HeLa-shNPC1 cell lysates treated with control siRNA (siScr) or siHDAC7. **C.** Bar graph depicting the ratio of total NPC1 to tubulin (left) and HDAC7 to tubulin (right). The NPC1 data is shown as the mean ± SD of the ratio of NPC1 to tubulin, whereas the HDAC7 data is shown as a normalized mean ± SD of the ratio of HDAC7 to tubulin with the siScr condition being set to 1. The asterisks represent p < 0.05 as determined by two-tailed t-test using siScr as the reference (n = 3). **D.** Representative Western blot of NPC1, HDAC7 and Tubulin from I1061T-expressing HeLa-shNPC1 cell lysates treated with control siRNA (siScr) or siHDAC7 in the absence (DMSO) or presence of 4mM VPA or 50 μM CQ for 24 h. **E.** Bar graph depicting the amount of EndoH^S^ (white) and EndoH^R^ (black) glycoforms as a fraction of total NPC1 from I1061T-expressing HeLa-shNPC1 cell lysates treated with control siRNA (siScr) or siHDAC7 in the absence (DMSO) or presence of 4mM VPA or 50 μM CQ for 24 h. The data are presented as the normalized mean ± SD and the asterisk indicates p < 0.05 using a two-tailed T-test with DMSO + siScr treatment as a reference (n = 3).

## Discussion

The HDACi VPA is widely used in the treatment of many neurological conditions including epilepsy and bipolar disorder due to its ability to cross the BBB and its minimal toxicity during chronic administration in mouse models and in humans (113,114). VPA has also exhibited therapeutic potential in mouse models of Alzheimer’s disease (AD) (115) and amyotrophic lateral sclerosis (ALS) (116), suggesting that it is capable of modulating the folding and/or aggregation of disease-associated variant proteins. This hypothesis is supported by our recent observations that HDACis are able to correct the trafficking and functional defects associated with disease-causing variants of CFTR and AAT (18,71,81,117-119).

Herein, we observed that VPA improves the trafficking and function of I1061T-NPC1, the most common NPC-associated variant, leading to restoration of cholesterol homeostasis in patient derived fibroblasts. This corrective property is dependent on the HDAC inhibitory activity of VPA, since its carboxamide derivative, valpromide, which lacks HDACi activity, is unable to correct the trafficking of I1061T-NPC1. While we did observe a VPA-mediated correction of the trafficking defect associated with the I1061T variant, it failed to exhibit increased stability in the LE/Ly compartment, thereby impacting its functional benefits. HDACis have been shown to alter the intracellular pH in the lysosome (96), thereby affecting compartment specific events which are likely to alter the stability and function of lysosomal proteins, such as NPC1. This effect can be overcome by a combinatorial treatment with another FDA-approved compound, chloroquine, which has been used as an anti-malarial agent for many years. CQ is a well-established lysosomotropic compound, which, in its free base form, can easily traverse the lipid bilayer of the lysosome and become ionized in its lumen, thereby reducing its permeability (120). The accumulation of CQ in the lumen of the lysosome serves as a sink for free protons culminating in an increase in the lysosomal pH (121), an effect which works to normalize the hyperacidification of the lysosome in response to HDACi treatment.

It is now accepted that HDACis alter the acetylation equilibrium of both histone (122) and non-histone proteins (45,123), which include numerous proteostasis components regulating the heat shock response (HSR) and unfolded protein response (UPR) programs (57,124), which are responsible for the recovery from protein folding stress such as the maladaptive stress response (MSR) caused by the chronic expression of misfolded variants of CFTR, AAT and NPC1 (119). Transcriptionally, acetylation of histones alters their ability to interact with DNA leading to altered expression profiles of many genes. HDACis promote the hyperacetylation of histones, thereby opening the chromatin structure leading to a global upregulation of gene expression as transcription factors have less restricted access to promoters. However, HDACis also regulate the acetylation of non-histone proteins (111), which can impede the ubiquitination of lysine residues, thereby altering protein stability, and modulating the activity of protein products by altering their ability to interact with substrates and regulatory proteins (125). Specifically, VPA has been shown to increase the activity of GRP78/Bip and Hsp70, cellular chaperones engaged in the folding and trafficking of *de novo* synthesized proteins (126,127), thereby providing protection from ER misfolding stress, which has been shown to occur in response to the chronic expression of I1061T-NPC1 (119).

While VPA can modulate the expression of numerous proteostasis components to promote the folding and trafficking of a functional I1061T-NPC1 polypeptide, we now show that it can promote the hyperacetylation of NPC1 itself. In order to understand the impact of NPC1 acetylation, we used the PAIL algorithm to identify likely sites of lysine acetylation in the NPC1 polypeptide. While we identified 16 lysine residues with high acetylation scores, we observed that only the loss of 4 of these lysine residues, K877, K1013, K1057 and K1217, impeded the ER export of WT-NPC1. Interestingly, 3 of these lysine residues are located in the CRD of NPC1, which is also the location of the I1061T mutation. These data suggest that the I1061T variant alters the folding of NPC1 in a way that limits the acetylation of the folded protein at one or multiple of these lysine residues, thereby contributing to the ER retention of the polypeptide. VPA, is able to restore the hyperacetylation of NPC1, thereby likely working in concert with the altered proteostasis environment to promote the trafficking of this disease-associated variant. While more work is required to fully elucidate the cohort of proteostasis changes associated with VPA treatment and to characterize their functional role in the biogenesis of NPC1, these data do suggest that VPA is able to provide a proteostasis environment amenable to the trafficking of a functional I1061T-NPC1 protein to the LE/Ly compartment.

While HDACis have been shown to have positive effects on the folding, trafficking and function of NPC1 (18,60,128), they have also been linked to restoring WT-like expression profiles for lipid biosynthetic components, including enzymes involved in cholesterol biosynthesis (129-131), which are dysregulated in NPC1 disease (132). Most notably, SAHA has been shown to normalize the NPC-associated elevation in gene expression related to cholesterol biosynthesis, including the sterol regulatory element binding factor 2 (*SREB2*), a transcription factor that regulates cholesterol biosynthesis components, as well as the expression of *Hmgcs1, Hmgcr, Mvk, Mvd, Idi1, Lss and Cyp51* (130). These data suggest that the therapeutic benefit of HDACis could be multifaceted, restoring balance to the dynamics of specialized compartments defining all steps of the endomembrane trafficking system. They are able to not only correct the trafficking defect associated with NPC1-linked variants to restore a functional NPC1 protein in the LE/Ly compartment, but they are also able to manage the dysregulated transcriptional profile of genes associated with lipid biosynthetic pathways, thereby abrogating lipotoxicity found in the brain and liver of NPC1 patients with advanced disease. While much work remains to address if VPA exhibits these latter properties in NPC1 disease, our data herein demonstrates that it is capable of correcting the trafficking defect linked with NPC1 variants.

The synergistic effect of complimentary compounds, such as observed with VPA and CQ could provide the opportunity to improve the therapeutic potential of cyclodextrin-based therapies. This hypothesis is supported by the report that the chronic administration of a triple combination formulation (TCF) of HPβCD/SAHA/PEG400 in the nmf164 (D1005G) mouse model (D1005G) provides improved benefit to disease progression (61). While this result has been challenged by the observation that SAHA is dispensible in TCF-mediated restoration of lipid homeostasis in a different, I1061T mouse model (62), our observations that a combinatorial treatment of cyclodextrins and VPA negatively impact the HDACi-mediated correction of I1061T-NPC1 trafficking suggest that drug-drug interactions or countering effects on critical cellular pathways could account for this observation. In light of these results, the proper course of action could require a sequential rather than combinatorial treatments with HDACi to achieve benefit.

While VPA is currently used to treat seizures in patients with late stage NPC1 disease, it had not been studied for its ability to abrogate the trafficking and lipid homeostasis defects linked to disease-causing NPC1 variants. We now provide evidence that VPA is able to correct NPC1 trafficking and cholesterol homeostasis using both human I1061T-expressing patient-derived fibroblasts and Hela cells, indicating its versatility in diverse human cellular environments. Of particular relevance to the VPA-mediated effects on NPC1 is the recent report that the monthly intra-thecal HPβCD dosing causes a statistically significant slowing of neuropsychological outcomes in NPC1 patients after 36 months (87). These findings suggest that the initial failings of VTS-270 to meet clinically significant benefit after 18 months (http://www.pharmatimes.com/news/mallinckrodts_niemann-pick_type_c_drug_fails_to_hit_trial_target_1259268) could simply be due to the shorter time course of the trial. Our results herein raise the possibility for the need of a combinatorial or sequential treatment approach to increase efficacy of HPβCD by increasing the functional pool of LE/Ly localized NPC1 variants undergoing HPβCD therapy which could accelerate the therapeutic window for patient benefit. It is possible that a combinatorial or sequential dosing approach using HPβCD, VPA and CQ could begin to more effectively address the complex genetic etiology of disease (1) to provide improved therapeutic benefit for NPC1 patients.

### Experimental Procedures

#### Reagents and Antibodies

Filipin, Valproic acid sodium salt, Chloroquine diphosphate salt, Methyl-β-cylodextrin (MβCD) and Acridine Orange hemi (zinc chloride) salt were purchased from Sigma-Aldrich (St. Louis, MO). Valpromide was purchased from Alfa Aesar (Ward Hill, MA). All restriction enzymes and endoglycosidase H (EndoH) were purchased from New England Biolabs (Ipswich, MA). Rat monoclonal anti-NPC1 antibody was developed against the C-terminal peptide (amino acids 1261-1278) of NPC1 protein. Rabbit polyclonal anti-NPC1 antibody was generously provided by Dan Ory (Washington University School of Medicine, St. Louis, MO), mouse anti-Tubulin (Sigma), rabbit anti-Calnexin (Stressgen), mouse anti-LAMP1 (obtained from the Developmental studies Hybridoma Bank). Rabbit anti-Alexa-488- and mouse anti-Alexa-647-conjugated secondary antibodies were obtained from Invitrogen (Carlsbad, CA). Cys/Met (EasyTag Express labeling kit) was purchased from PerkinElmer Life Sciences.

#### Plasmids

cDNA encoding human ΔU3m*npc1*-WT construct was kindly provided by Dan Ory (133). *npc1* gene was subcloned into pLVX-IRES-Neo (Clontech) lentiviral vector using EcoR1 and BamH1 cloning sites. Mission shRNA clone was purchased from sigma against 3’UTR (TRCN000000542) of human NPC1 (pLKO-shNPC1) for knockdown of *npc1* gene in human cell lines. GFP-KDEL construct was generated by inserting calreticulin signal peptide (1-18 amino acid) sequences at the N-terminal of GFP in pEGFP-N1 vector (Clontech) and the ER retention signal KDEL sequence was incorporated at the C-terminal region of GFP with stop codon by seamless cloning technique (Stratagene, La Jolla). Using human *TMEM97* cDNA (Open Biosystems) as template, TMEM97 gene was integrated into pEGFP-N1 (Clontech) plasmid at BamH1, EcoR1 restriction sites. All the human NPC1 (WT and I1061T) and other constructs used in this study was confirmed by DNA sequencing.

#### Cell lines, Cell culture

Human wild-type fibroblasts (GM05659), and NPC1 mutant fibroblasts; I1061T/I1061T (GM18453), P237S/I1061T (GM3123) were purchased from Coriell Cell Repositories (Coriell institute for Medical Research). Human fibroblast cells were grown in DMEM medium supplemented with 2 mM L-Glutamine, 10% FBS, 50-units/ml penicillin and 50 μg/ml streptomycin antibiotics. The endogenous *npc1* gene in HeLa cells was silenced with 3’-UTR shNPC1 lentivirus (sigma) to generate the *npc1-*deficient cells and stable clones were selected with 3 μg/ml puromycin antibiotics for two weeks. HeLa-shNPC1 cells stably expressing human NPC1 WT or I1061T mutant was generated using lentiviral construct (pLVX-Neo) and selection was done with 600 μg/ml G418 antibiotics. Stable clones expressing of WT or I1061T NPC1 mutant protein were cultured in DMEM medium supplemented with 10% FBS, 50-units/ml penicillin, 50 μg/ml streptomycin, 3 μg/ml puromycin, and 600 μg/ml G418 antibiotics.

#### Cell transfection and lentivirus transduction

GFP-KDEL or TMEM97-GFP plasmids were transfected in HeLa-shNPC1 cells stably expressing NPC1-I1061T mutant using Lipofectamine LTX or Lipofectamine 2000 (Invitrogen) reagents. siRNA targeting against HDACs 1-11 were transfected using Lipofectamine RNAiMAX according to the instructions provided by the manufacture (Invitrogen). Stable expression of human NPC1 WT and I1061T mutant in HeLa-shNPC1 cells was done with lentivirus expression system. Briefly, HEK293T cells were seeded and grown to 60-70% confluence in 25 cm dish. Cells were transfected with pLVX-Neo plasmid expressing human NPC1 WT or I1061T mutant along with the lentivirus helper plasmids RRE, REV, VSVG in Opti-MEM medium using Fugene 6 transfection reagents (Promega). Lentiviruses were collected from HEK293T cells after 36 and 48 h post-transfection. Spun the collected virus at 1800 rpm to get rid of the cell debris. The lentiviral particles were filtered through 0.45 μM filters (Millipore), followed by high-speed centrifugation at 50,000 x g at RT for 2 h. The translucent lentivirus pellet was resuspended in 200 μl HBSS (14175-095 Invitrogen) supplemented with 0.5 mM MgCl_2_, 0.4 mM MgSO_4_, 1 mM CaCl_2_. Small aliquots of the concentrated lentiviruses were stored at −80 °C.

#### Endoglycosidase H assay

Human NPC1 WT and I1061T mutant stably expressed in HeLa-shNPC1 cells were washed with 1x PBS twice and lysed the cells with RIPA lysis buffer (10 mM Tris pH 8.0, 140 mM NaCl, 1% NP-40, 0.1% sodium-deoxycholate, 0.1% SDS, 1x protease inhibitor cocktail (PIC), 1 mM PMSF) on ice for 30 min. Cell lysates were obtained by 16,000 x g centrifugation at 4 °C. Protein concentration was measured using standard BCA assay. ∼300 μg of cell lysates were incubated with 1-2 μg of rabbit polyclonal anti-NPC1 antibody or rat monoclonal anti-NPC1 antibody overnight at 4°C. 20 μl of protein A/G-Sepharose or GammaBind G Sepharose beads (GE Healthcare) were added to the cell lysates to capture the antibody bound NPC1 and incubated for 2 h at 4°C. The beads were washed with RIPA lysis buffer three times and 1 x with PBS. The immunoprecipitated NPC1 was eluted with 1 x denaturation buffer (0.5% SDS, 40 mM DTT) at 95°C for 10min. Immunoprecipitated NPC1 was divided into equal parts and incubated in the presence (+) or absence (-) of 1,000 units Endoglycosidase H (EndoH) enzyme overnight at 37°C. EndoH digested samples were subjected to 4-12% Bis-tris gradient (Invitrogen) or 4-20% gradient (Bio-Rad) SDS-PAGE gels and immunoblotted with anti-NPC1 antibody (1: 3000).

### qRT-PCR

Total RNA was isolated from cells using TRIzol reagent (Invitrogen) according to the manufactures instructions. cDNA was generated for 10 min at 50°C then target products amplified 40 cycles in the presence of SYBR Green PCR master mixture (Quanta Biosciences) using template-specific primers (200 nm) in a DNA Engine Opticon 2 Real-Time cycler system (Bio-Rad). The following primer sequences were used for the qRT-PCR: human *NPC1* (forward, 5′-gtctccgagtacactcccatc-3′; reverse, 5′-cgcagtaatgaagaccagcga-3′) as described in (134) and human glyceraldehyde-3-phosphate dehydrogenase (forward, 5′-gagtcaacggattggtcgt-3’; reverse, 5′-gaggtcaatgaaggggtcat-3′). Relative quantification of gene expression was performed using the comparative threshold (*C*_T_) method as described by the manufacturer. Relative changes in mRNA expression level were calculated following normalization to glyceraldehyde-3-phosphate dehydrogenase expression based on three independent assays.

### Metabolic labeling of NPC1 protein

Pulse-chase of human NPC1 protein was performed by incubating the HeLa-shNPC1 cells stably expressing WT- or I1061T-NPC1 for 1h in starvation media devoid of Cys/Met, followed by a 1h incubation in labeling media supplemented with 100μCi [^35^ S]-labeled Cys and Met (EasyTag Express labeling kit, PerkinElmer). After the 1h pulse, cells were chased for the indicated time, washed with twice with PBS, followed by lysis, as described for SDS-PAGE sample preparation above, for 30min on ice. Cell lysates were collected at 14,000xrpm for 10min and protein concentration was measured by standard BCA assay. Equal amount of cell lysate was incubated with NPC1 antibody cross-linked to GammaBinding Sepharose beads and incubated overnight at 4°C. NPC1 protein was eluted as described above and EndoH treatment was performed as described above. NPC1 proteins were separated in 4-20% Bio-Rad gradient gels and analyzed by autoradiography.

### Microscopy

For immunofluorescence cells were grown on the 22-mm coverslip in 6-well plate. Cells were washed twice with PBS, fixed with fresh 4% formaldehyde in PBS for 10min. Cells were subsequently washed 3 times with PBS containing 1.5 mg/ml glycine to quench any remaining formaldehyde. Cells were subsequently blocked with 10% goat serum in PBS and stained with 25 μg/ml filipin for 30 min at RT. Cells were washed twice with PBS and probed with rabbit anti-NPC1 (1:1000) and mouse anti-LAMP1 (1:1000) for 2 h at RT. Cells were washed three times with PBS, incubated with Alexa Flour secondary antibodies (Invitrogen), goat anti-rabbit Alexa-488 (1:1000) and goat anti-mouse Alexa-647 (1:1000) for 1 h, washed 3 times with PBS, coverslips were mounted using Fluoromount G solution (Electron Microscopy Sciences) on the glass slide. Quantitative filipin staining was performed as previously described (92) and the images were taken using an A4 filter cube (Leica, Wetzlar, Germany). Fluorescence crossover from one channel to another was measured using single-labeled samples of each probe and found to be insignificant. Images were background-corrected as previously described (135). To analyze the lysosomal pH and integrity of lysosomal membranes, cells were treated with drugs as described in *Results* for 48 h and then incubated with acridine orange (2 μg/ml) for 30 min at 37 °C, washed several times with 1x PBS, mounted the coverslips on the glass slide. Cells were visualized with confocal microscopy with the appropriate laser settings of excitation at 460 nm and emission at 620 nm.

### Statistical analysis

All statistical analyses were performed with the assistance of GraphPad prism software (GraphPad Software, San Diego, CA).

## Acknowledgements

We thank Daniel Ory (Washington University, St. Louis, MO) for providing polyclonal anti-rabbit NPC1 antibody. We thank Frederic Maxfield (Weill Cornell Medical College, New York, NY) for his valuable comments and technical expertise is gathering and analyzing free cholesterol images. We thank the Support for Accelerated Research for Nieman-Pick Type C (SOAR) and the Ara Parseghian Medical Research Fund (APMRF), University of Notre Dame for postdoctoral support for Kanagarj Subramanian. The LAMP1 antibody 1D4B was obtained from the Developmental Studies Hybridoma Bank, created by the NICHD of the NIH, and maintained at The University of Iowa. This work is supported by NIH grants AG049665 and HL095524 to WEB.

## References

1. Wang, C., and Balch, W. E. (2018) Bridging Genomics to Phenomics at Atomic Resolution through Variation Spatial Profiling. Cell Rep 24, 2013–2028 e2016

2. Fog, C. K., and Kirkegaard, T. (2019) Animal models for Niemann-Pick type C: implications for drug discovery & development. Expert Opin Drug Discov 14, 499–509

3. Torres, S., Balboa, E., Zanlungo, S., Enrich, C., Garcia-Ruiz, C., and Fernandez-Checa, J. C. (2017) Lysosomal and Mitochondrial Liaisons in Niemann-Pick Disease. Front Physiol 8, 982

4. Ordonez, M. P., and Steele, J. W. (2017) Modeling Niemann Pick type C1 using human embryonic and induced pluripotent stem cells. Brain Res 1656, 63–67

5. Vanier, M. T. (2015) Complex lipid trafficking in Niemann-Pick disease type C. J Inherit Metab Dis 38, 187–199

6. Vivas, O., Tiscione, S. A., Dixon, R. E., Ory, D. S., and Dickson, E. J. (2019) Niemann-Pick Type C Disease Reveals a Link between Lysosomal Cholesterol and PtdIns(4,5)P2 That Regulates Neuronal Excitability. Cell Rep 27, 2636–2648 e2634

7. Pugach, E. K., Feltes, M., Kaufman, R. J., Ory, D. S., and Bang, A. G. (2018) High-content screen for modifiers of Niemann-Pick type C disease in patient cells. Hum Mol Genet 27, 2101–2112

8. Ory, D. S. (2004) The niemann-pick disease genes; regulators of cellular cholesterol homeostasis. Trends Cardiovasc Med 14, 66–72

9. Frolov, A., Srivastava, K., Daphna-Iken, D., Traub, L. M., Schaffer, J. E., and Ory, D. S. (2001) Cholesterol overload promotes morphogenesis of a Niemann-Pick C (NPC)-like compartment independent of inhibition of NPC1 or HE1/NPC2 function. J Biol Chem 276, 46414–46421

10. Zhang, M., Dwyer, N. K., Love, D. C., Cooney, A., Comly, M., Neufeld, E., Pentchev, P. G., Blanchette-Mackie, E. J., and Hanover, J. A. (2001) Cessation of rapid late endosomal tubulovesicular trafficking in Niemann-Pick type C1 disease. Proceedings of the National Academy of Sciences of the United States of America 98, 4466–4471

11. Neufeld, E. B., Cooney, A. M., Pitha, J., Dawidowicz, E. A., Dwyer, N. K., Pentchev, P. G., and Blanchette-Mackie, E. J. (1996) Intracellular trafficking of cholesterol monitored with a cyclodextrin. J Biol Chem 271, 21604–21613

12. Poirier, S., Mayer, G., Murphy, S. R., Garver, W. S., Chang, T. Y., Schu, P., and Seidah, N. G. (2013) The cytosolic adaptor AP-1A is essential for the trafficking and function of Niemann-Pick type C proteins. Traffic 14, 458–469

13. Kwon, H. J., Abi-Mosleh, L., Wang, M. L., Deisenhofer, J., Goldstein, J. L., Brown, M. S., and Infante, R. E. (2009) Structure of N-terminal domain of NPC1 reveals distinct subdomains for binding and transfer of cholesterol. Cell 137, 1213–1224

14. Deffieu, M. S., and Pfeffer, S. R. (2011) Niemann-Pick type C 1 function requires lumenal domain residues that mediate cholesterol-dependent NPC2 binding. Proceedings of the National Academy of Sciences of the United States of America 108, 18932–18936

15. Estiu, G., Khatri, N., and Wiest, O. (2013) Computational studies of the cholesterol transport between NPC2 and the N-terminal domain of NPC1 (NPC1(NTD)). Biochemistry 52, 6879–6891

16. Hindorff, L. A., Bonham, V. L., Brody, L. C., Ginoza, M. E. C., Hutter, C. M., Manolio, T. A., and Green, E. D. (2018) Prioritizing diversity in human genomics research. Nat Rev Genet 19, 175–185

17. Wang, C., Scott, S. M., Hutt, D. M., Zhao, P., Shao, H., Gestwicki, J. E., and Balch, W. E. (2018) Managing the spatial covariance of genetic diversity in Niemann-Pick C1 through modulation of the Hsp70 chaperone system. bioRxiv doi: http://dx.doi.org/10.1101/437764.

18. Pipalia, N. H., Subramanian, K., Mao, S., Ralph, H., Hutt, D. M., Scott, S. M., Balch, W. E., and Maxfield, F. R. (2017) Histone deacetylase inhibitors correct the cholesterol storage defect in most Niemann-Pick C1 mutant cells. J Lipid Res 58, 695–708

19. Elrick, M. J., Pacheco, C. D., Yu, T., Dadgar, N., Shakkottai, V. G., Ware, C., Paulson, H. L., and Lieberman, A. P. (2010) Conditional Niemann-Pick C mice demonstrate cell autonomous Purkinje cell neurodegeneration. Hum Mol Genet 19, 837–847

20. Totenhagen, J. W., Bernstein, A., Yoshimaru, E. S., Erickson, R. P., and Trouard, T. P. (2017) Quantitative magnetic resonance imaging of brain atrophy in a mouse model of Niemann-Pick type C disease. PLoS One 12, e0178179

21. Totenhagen, J. W., Lope-Piedrafita, S., Borbon, I. A., Yoshimaru, E. S., Erickson, R. P., and Trouard, T. P. (2012) In vivo assessment of neurodegeneration in Niemann-Pick type C mice by quantitative T2 mapping and diffusion tensor imaging. J Magn Reson Imaging 35, 528–536

22. Totenhagen, J. W., Yoshimaru, E. S., Erickson, R. P., and Trouard, T. P. (2013) (1) H magnetic resonance spectroscopy of neurodegeneration in a mouse model of niemann-pick type C1 disease. J Magn Reson Imaging 37, 1195–1201

23. Zaaraoui, W., Crespy, L., Rico, A., Faivre, A., Soulier, E., Confort-Gouny, S., Cozzone, P. J., Pelletier, J., Ranjeva, J. P., Kaphan, E., and Audoin, B. (2011) In vivo quantification of brain injury in adult Niemann-Pick Disease Type C. Mol Genet Metab 103, 138–141

24. Millat, G., Marcais, C., Rafi, M. A., Yamamoto, T., Morris, J. A., Pentchev, P. G., Ohno, K., Wenger, D. A., and Vanier, M. T. (1999) Niemann-Pick C1 disease: the I1061T substitution is a frequent mutant allele in patients of Western European descent and correlates with a classic juvenile phenotype. Am J Hum Genet 65, 1321–1329

25. Park, W. D., O’Brien, J. F., Lundquist, P. A., Kraft, D. L., Vockley, C. W., Karnes, P. S., Patterson, M. C., and Snow, K. (2003) Identification of 58 novel mutations in Niemann-Pick disease type C: correlation with biochemical phenotype and importance of PTC1-like domains in NPC1. Hum Mutat 22, 313–325

26. Vanier, M. T., and Millat, G. (2003) Niemann-Pick disease type C. Clinical genetics 64, 269–281

27. Ko, D. C., Milenkovic, L., Beier, S. M., Manuel, H., Buchanan, J., and Scott, M. P. (2005) Cell-autonomous death of cerebellar purkinje neurons with autophagy in Niemann-Pick type C disease. PLoS genetics 1, 81–95

28. Mengel, E., Klunemann, H. H., Lourenco, C. M., Hendriksz, C. J., Sedel, F., Walterfang, M., and Kolb, S. A. (2013) Niemann-Pick disease type C symptomatology: an expert-based clinical description. Orphanet J Rare Dis 8, 166

29. Runz, H., Dolle, D., Schlitter, A. M., and Zschocke, J. (2008) NPC-db, a Niemann-Pick type C disease gene variation database. Hum Mutat 29, 345–350

30. Scott, C., and Ioannou, Y. A. (2004) The NPC1 protein: structure implies function. Biochim Biophys Acta 1685, 8–13

31. Millat, G., Marcais, C., Tomasetto, C., Chikh, K., Fensom, A. H., Harzer, K., Wenger, D. A., Ohno, K., and Vanier, M. T. (2001) Niemann-Pick C1 disease: correlations between NPC1 mutations, levels of NPC1 protein, and phenotypes emphasize the functional significance of the putative sterol-sensing domain and of the cysteine-rich luminal loop. Am J Hum Genet 68, 1373–1385

32. Zech, M., Nubling, G., Castrop, F., Jochim, A., Schulte, E. C., Mollenhauer, B., Lichtner, P., Peters, A., Gieger, C., Marquardt, T., Vanier, M. T., Latour, P., Klunemann, H., Trenkwalder, C., Diehl-Schmid, J., Perneczky, R., Meitinger, T., Oexle, K., Haslinger, B., Lorenzl, S., and Winkelmann, J. (2013) Niemann-Pick C disease gene mutations and age-related neurodegenerative disorders. PLoS One 8, e82879

33. Gelsthorpe, M. E., Baumann, N., Millard, E., Gale, S. E., Langmade, S. J., Schaffer, J. E., and Ory, D. S. (2008) Niemann-Pick type C1 I1061T mutant encodes a functional protein that is selected for endoplasmic reticulum-associated degradation due to protein misfolding. J Biol Chem 283, 8229–8236

34. Fog, C. K., Zago, P., Malini, E., Solanko, L. M., Peruzzo, P., Bornaes, C., Magnoni, R., Mehmedbasic, A., Petersen, N. H. T., Bembi, B., Aerts, J., Dardis, A., and Kirkegaard, T. (2018) The heat shock protein amplifier arimoclomol improves refolding, maturation and lysosomal activity of glucocerebrosidase. EBioMedicine 38, 142–153

35. Kirkegaard, T., Gray, J., Priestman, D. A., Wallom, K. L., Atkins, J., Olsen, O. D., Klein, A., Drndarski, S., Petersen, N. H., Ingemann, L., Smith, D. A., Morris, L., Bornaes, C., Jorgensen, S. H., Williams, I., Hinsby, A., Arenz, C., Begley, D., Jaattela, M., and Platt, F. M. (2016) Heat shock protein-based therapy as a potential candidate for treating the sphingolipidoses. Sci Transl Med 8, 355ra118

36. Deane, C. A., and Brown, I. R. (2016) Induction of heat shock proteins in differentiated human neuronal cells following co-application of celastrol and arimoclomol. Cell Stress Chaperones 21, 837–848

37. Parfitt, D. A., Aguila, M., McCulley, C. H., Bevilacqua, D., Mendes, H. F., Athanasiou, D., Novoselov, S. S., Kanuga, N., Munro, P. M., Coffey, P. J., Kalmar, B., Greensmith, L., and Cheetham, M. E. (2014) The heat-shock response co-inducer arimoclomol protects against retinal degeneration in rhodopsin retinitis pigmentosa. Cell Death Dis 5, e1236

38. Camargo, F., Erickson, R. P., Garver, W. S., Hossain, G. S., Carbone, P. N., Heidenreich, R. A., and Blanchard, J. (2001) Cyclodextrins in the treatment of a mouse model of Niemann-Pick C disease. Life Sci 70, 131–142

39. Davidson, C. D., Ali, N. F., Micsenyi, M. C., Stephney, G., Renault, S., Dobrenis, K., Ory, D. S., Vanier, M. T., and Walkley, S. U. (2009) Chronic cyclodextrin treatment of murine Niemann-Pick C disease ameliorates neuronal cholesterol and glycosphingolipid storage and disease progression. PLoS One 4, e6951

40. Liu, B., Turley, S. D., Burns, D. K., Miller, A. M., Repa, J. J., and Dietschy, J. M. (2009) Reversal of defective lysosomal transport in NPC disease ameliorates liver dysfunction and neurodegeneration in the npc1-/-mouse. Proceedings of the National Academy of Sciences of the United States of America 106, 2377–2382

41. Vite, C. H., Bagel, J. H., Swain, G. P., Prociuk, M., Sikora, T. U., Stein, V. M., O’Donnell, P., Ruane, T., Ward, S., Crooks, A., Li, S., Mauldin, E., Stellar, S., De Meulder, M., Kao, M. L., Ory, D. S., Davidson, C., Vanier, M. T., and Walkley, S. U. (2015) Intracisternal cyclodextrin prevents cerebellar dysfunction and Purkinje cell death in feline Niemann-Pick type C1 disease. Sci Transl Med 7, 276ra226

42. Balasubramanyam, K., Altaf, M., Varier, R. A., Swaminathan, V., Ravindran, A., Sadhale, P. P., and Kundu, T. K. (2004) Polyisoprenylated benzophenone, garcinol, a natural histone acetyltransferase inhibitor, represses chromatin transcription and alters global gene expression. J Biol Chem 279, 33716–33726

43. Bowers, E. M., Yan, G., Mukherjee, C., Orry, A., Wang, L., Holbert, M. A., Crump, N. T., Hazzalin, C. A., Liszczak, G., Yuan, H., Larocca, C., Saldanha, S. A., Abagyan, R., Sun, Y., Meyers, D. J., Marmorstein, R., Mahadevan, L. C., Alani, R. M., and Cole, P. A. (2010) Virtual ligand screening of the p300/CBP histone acetyltransferase: identification of a selective small molecule inhibitor. Chem Biol 17, 471–482

44. Lau, O. D., Kundu, T. K., Soccio, R. E., Ait-Si-Ali, S., Khalil, E. M., Vassilev, A., Wolffe, A. P., Nakatani, Y., Roeder, R. G., and Cole, P. A. (2000) HATs off: selective synthetic inhibitors of the histone acetyltransferases p300 and PCAF. Mol Cell 5, 589–595

45. Choudhary, C., Kumar, C., Gnad, F., Nielsen, M. L., Rehman, M., Walther, T. C., Olsen, J. V., and Mann, M. (2009) Lysine acetylation targets protein complexes and co-regulates major cellular functions. Science 325, 834–840

46. Peserico, A., and Simone, C. (2011) Physical and functional HAT/HDAC interplay regulates protein acetylation balance. J Biomed Biotechnol 2011, 371832

47. Marks, P. A., and Breslow, R. (2007) Dimethyl sulfoxide to vorinostat: development of this histone deacetylase inhibitor as an anticancer drug. Nat Biotechnol 25, 84–90

48. Garnock-Jones, K. P. (2015) Panobinostat: first global approval. Drugs 75, 695–704

49. VanderMolen, K. M., McCulloch, W., Pearce, C. J., and Oberlies, N. H. (2011) Romidepsin (Istodax, NSC 630176, FR901228, FK228, depsipeptide): a natural product recently approved for cutaneous T-cell lymphoma. J Antibiot (Tokyo) 64, 525–531

50. Milne, J. C., and Denu, J. M. (2008) The Sirtuin family: therapeutic targets to treat diseases of aging. Curr Opin Chem Biol 12, 11–17

51. Akerfelt, M., Morimoto, R. I., and Sistonen, L. (2010) Heat shock factors: integrators of cell stress, development and lifespan. Nat Rev Mol Cell Biol 11, 545–555

52. Scroggins, B. T., Robzyk, K., Wang, D., Marcu, M. G., Tsutsumi, S., Beebe, K., Cotter, R. J., Felts, S., Toft, D., Karnitz, L., Rosen, N., and Neckers, L. (2007) An acetylation site in the middle domain of Hsp90 regulates chaperone function. Mol Cell 25, 151–159

53. Nguyen, M. T., Somogyvari, M., and Soti, C. (2018) Hsp90 Stabilizes SIRT1 Orthologs in Mammalian Cells and C. elegans. Int J Mol Sci 19

54. Verma, P., Pfister, J. A., Mallick, S., and D’Mello, S. R. (2014) HSF1 protects neurons through a novel trimerization- and HSP-independent mechanism. J Neurosci 34, 1599–1612

55. Grandjean, J. M. D., Plate, L., Morimoto, R. I., Bollong, M. J., Powers, E. T., and Wiseman, R. L. (2019) Deconvoluting Stress-Responsive Proteostasis Signaling Pathways for Pharmacologic Activation Using Targeted RNA Sequencing. ACS Chem Biol 14, 784–795

56. Li, J., Labbadia, J., and Morimoto, R. I. (2017) Rethinking HSF1 in Stress, Development, and Organismal Health. Trends Cell Biol 27, 895–905

57. Rao, R., Fiskus, W., Ganguly, S., Kambhampati, S., and Bhalla, K. N. (2012) HDAC inhibitors and chaperone function. Adv Cancer Res 116, 239–262

58. Suraweera, A., O’Byrne, K. J., and Richard, D. J. (2018) Combination Therapy With Histone Deacetylase Inhibitors (HDACi) for the Treatment of Cancer: Achieving the Full Therapeutic Potential of HDACi. Front Oncol 8, 92

59. Munkacsi, A. B., Chen, F. W., Brinkman, M. A., Higaki, K., Gutierrez, G. D., Chaudhari, J., Layer, J. V., Tong, A., Bard, M., Boone, C., Ioannou, Y. A., and Sturley, S. L. (2011) An “exacerbate-reverse” strategy in yeast identifies histone deacetylase inhibition as a correction for cholesterol and sphingolipid transport defects in human Niemann-Pick type C disease. J Biol Chem 286, 23842–23851

60. Pipalia, N. H., Cosner, C. C., Huang, A., Chatterjee, A., Bourbon, P., Farley, N., Helquist, P., Wiest, O., and Maxfield, F. R. (2011) Histone deacetylase inhibitor treatment dramatically reduces cholesterol accumulation in Niemann-Pick type C1 mutant human fibroblasts. Proceedings of the National Academy of Sciences of the United States of America 108, 5620–5625

61. Alam, M. S., Getz, M., and Haldar, K. (2016) Chronic administration of an HDAC inhibitor treats both neurological and systemic Niemann-Pick type C disease in a mouse model. Sci Transl Med 8, 326ra323

62. Davidson, J., Molitor, E., Moores, S., Gale, S. E., Subramanian, K., Jiang, X., Sidhu, R., Kell, P., Zhang, J., Fujiwara, H., Davidson, C., Helquist, P., Melancon, B. J., Grigalunas, M., Liu, G., Salahi, F., Wiest, O., Xu, X., Porter, F. D., Pipalia, N. H., Cruz, D. L., Holson, E. B., Schaffer, J. E., Walkley, S. U., Maxfield, F. R., and Ory, D. S. (2019) 2-Hydroxypropyl-beta-cyclodextrin is the active component in a triple combination formulation for treatment of Niemann-Pick C1 disease. Biochim Biophys Acta Mol Cell Biol Lipids

63. Christensen, D. P., Dahllof, M., Lundh, M., Rasmussen, D. N., Nielsen, M. D., Billestrup, N., Grunnet, L. G., and Mandrup-Poulsen, T. (2011) Histone deacetylase (HDAC) inhibition as a novel treatment for diabetes mellitus. Mol Med 17, 378–390

64. Ozcan, U., Yilmaz, E., Ozcan, L., Furuhashi, M., Vaillancourt, E., Smith, R. O., Gorgun, C. Z., and Hotamisligil, G. S. (2006) Chemical chaperones reduce ER stress and restore glucose homeostasis in a mouse model of type 2 diabetes. Science 313, 1137–1140

65. Jagannath, S., Dimopoulos, M. A., and Lonial, S. (2010) Combined proteasome and histone deacetylase inhibition: A promising synergy for patients with relapsed/refractory multiple myeloma. Leuk Res 34, 1111–1118

66. Leggatt, G. R., and Gabrielli, B. (2012) Histone deacetylase inhibitors in the generation of the anti-tumour immune response. Immunol Cell Biol 90, 33–38

67. Marks, P. A. (2010) The clinical development of histone deacetylase inhibitors as targeted anticancer drugs. Expert Opin Investig Drugs 19, 1049–1066

68. Buckland, J. (2011) Rheumatoid arthritis: HDAC and HDACi: pathogenetic and mechanistic insights. Nat Rev Rheumatol 7, 682

69. Wang, C., Bouchecareilh, M., and Balch, W. E. (2017) Measuring the Effect of Histone Deacetylase Inhibitors (HDACi) on the Secretion and Activity of Alpha-1 Antitrypsin. Methods Mol Biol 1639, 185–193

70. Roth, D. M., Hutt, D. M., Tong, J., Bouchecareilh, M., Wang, N., Seeley, T., Dekkers, J. F., Beekman, J. M., Garza, D., Drew, L., Masliah, E., Morimoto, R. I., and Balch, W. E. (2014) Modulation of the maladaptive stress response to manage diseases of protein folding. PLoS Biol 12, e1001998

71. Bouchecareilh, M., Hutt, D. M., Szajner, P., Flotte, T. R., and Balch, W. E. (2012) Histone deacetylase inhibitor (HDACi) suberoylanilide hydroxamic acid (SAHA)-mediated correction of alpha1-antitrypsin deficiency. J Biol Chem 287, 38265–38278

72. Bouchecareilh, M., and Balch, W. E. (2012) Proteostasis, an emerging therapeutic paradigm for managing inflammatory airway stress disease. Curr Mol Med 12, 815–826

73. van den Bosch, T., Kwiatkowski, M., Bischoff, R., and Dekker, F. J. (2017) Targeting transcription factor lysine acetylation in inflammatory airway diseases. Epigenomics 9, 1013–1028

74. Royce, S. G., Moodley, Y., and Samuel, C. S. (2014) Novel therapeutic strategies for lung disorders associated with airway remodelling and fibrosis. Pharmacol Ther 141, 250–260

75. Marwick, J. A., Adcock, I. M., and Chung, K. F. (2010) Overcoming reduced glucocorticoid sensitivity in airway disease: molecular mechanisms and therapeutic approaches. Drugs 70, 929–948

76. Marwick, J. A., Ito, K., Adcock, I. M., and Kirkham, P. A. (2007) Oxidative stress and steroid resistance in asthma and COPD: pharmacological manipulation of HDAC-2 as a therapeutic strategy. Expert Opin Ther Targets 11, 745–755

77. Halili, M. A., Andrews, M. R., Sweet, M. J., and Fairlie, D. P. (2009) Histone deacetylase inhibitors in inflammatory disease. Curr Top Med Chem 9, 309–319

78. Banerjee, A., Trivedi, C. M., Damera, G., Jiang, M., Jester, W., Hoshi, T., Epstein, J. A., and Panettieri, R. A., Jr. (2012) Trichostatin A abrogates airway constriction, but not inflammation, in murine and human asthma models. Am J Respir Cell Mol Biol 46, 132–138

79. Bouchecareilh, M., and Balch, W. E. (2011) Proteostasis: a new therapeutic paradigm for pulmonary disease. Proc Am Thorac Soc 8, 189–195

80. Angles, F., Hutt, D. M., and Balch, W. E. (2019) HDAC Inhibitors Rescue Multiple Disease-Causing CFTR Variants. Hum Mol Genet

81. Hutt, D. M., Herman, D., Rodrigues, A. P., Noel, S., Pilewski, J. M., Matteson, J., Hoch, B., Kellner, W., Kelly, J. W., Schmidt, A., Thomas, P. J., Matsumura, Y., Skach, W. R., Gentzsch, M., Riordan, J. R., Sorscher, E. J., Okiyoneda, T., Yates, J. R., 3rd, Lukacs, G. L., Frizzell, R. A., Manning, G., Gottesfeld, J. M., and Balch, W. E. (2010) Reduced histone deacetylase 7 activity restores function to misfolded CFTR in cystic fibrosis. Nature chemical biology 6, 25–33

82. Hutt, D. M., Olsen, C. A., Vickers, C. J., Herman, D., Chalfant, M., Montero, A., Leman, L. J., Burkle, R., Maryanoff, B. E., Balch, W. E., and Ghadiri, M. R. (2011) Potential Agents for Treating Cystic Fibrosis: Cyclic Tetrapeptides that Restore Trafficking and Activity of DeltaF508-CFTR. ACS Med Chem Lett 2, 703–707

83. Chiu, C. T., Wang, Z., Hunsberger, J. G., and Chuang, D. M. (2013) Therapeutic potential of mood stabilizers lithium and valproic acid: beyond bipolar disorder. Pharmacol Rev 65, 105–142

84. Hershey, L. A., and Coleman-Jackson, R. (2019) Pharmacological Management of Dementia with Lewy Bodies. Drugs Aging 36, 309–319

85. Singh, A. K., Halder-Sinha, S., Clement, J. P., and Kundu, T. K. (2018) Epigenetic modulation by small molecule compounds for neurodegenerative disorders. Pharmacol Res 132, 135–148

86. Ketter, T. A., Frye, M. A., Cora-Locatelli, G., Kimbrell, T. A., and Post, R. M. (1999) Metabolism and excretion of mood stabilizers and new anticonvulsants. Cell Mol Neurobiol 19, 511–532

87. Farmer, C. A., Thurm, A., Farhat, N., Bianconi, S., Keener, L. A., and Porter, F. D. (2019) Long-Term Neuropsychological Outcomes from an Open-Label Phase I/IIa Trial of 2-Hydroxypropyl-beta-Cyclodextrins (VTS-270) in Niemann-Pick Disease, Type C1. CNS Drugs 33, 677–683

88. Hussein, Z., Mukherjee, D., Lamm, J., Cavanaugh, J. H., and Granneman, G. R. (1994) Pharmacokinetics of valproate after multiple-dose oral and intravenous infusion administration: gastrointestinal-related diurnal variation. J Clin Pharmacol 34, 754–759

89. Slater, A. F. (1993) Chloroquine: mechanism of drug action and resistance in Plasmodium falciparum. Pharmacol Ther 57, 203–235

90. Dunmore, B. J., Drake, K. M., Upton, P. D., Toshner, M. R., Aldred, M. A., and Morrell, N. W. (2013) The lysosomal inhibitor, chloroquine, increases cell surface BMPR-II levels and restores BMP9 signalling in endothelial cells harbouring BMPR-II mutations. Hum Mol Genet 22, 3667–3679

91. Helquist, P., Maxfield, F. R., Wiech, N. L., and Wiest, O. (2013) Treatment of Niemann--pick type C disease by histone deacetylase inhibitors. Neurotherapeutics 10, 688–697

92. Pipalia, N. H., Huang, A., Ralph, H., Rujoi, M., and Maxfield, F. R. (2006) Automated microscopy screening for compounds that partially revert cholesterol accumulation in Niemann-Pick C cells. J Lipid Res 47, 284–301

93. Schultz, M. L., Krus, K. L., Kaushik, S., Dang, D., Chopra, R., Qi, L., Shakkottai, V. G., Cuervo, A. M., and Lieberman, A. P. (2018) Coordinate regulation of mutant NPC1 degradation by selective ER autophagy and MARCH6-dependent ERAD. Nat Commun 9, 3671

94. Davies, J. P., and Ioannou, Y. A. (2000) Topological analysis of Niemann-Pick C1 protein reveals that the membrane orientation of the putative sterol-sensing domain is identical to those of 3-hydroxy-3-methylglutaryl-CoA reductase and sterol regulatory element binding protein cleavage-activating protein. J Biol Chem 275, 24367–24374

95. Eriksson, I., Joosten, M., Roberg, K., and Ollinger, K. (2013) The histone deacetylase inhibitor trichostatin A reduces lysosomal pH and enhances cisplatin-induced apoptosis. Experimental cell research 319, 12–20

96. McBrian, M. A., Behbahan, I. S., Ferrari, R., Su, T., Huang, T. W., Li, K., Hong, C. S., Christofk, H. R., Vogelauer, M., Seligson, D. B., and Kurdistani, S. K. (2013) Histone acetylation regulates intracellular pH. Mol Cell 49, 310–321

97. Krogstad, D. J., and Schlesinger, P. H. (1987) The basis of antimalarial action: non-weak base effects of chloroquine on acid vesicle pH. Am J Trop Med Hyg 36, 213–220

98. Davies, J. P., Chen, F. W., and Ioannou, Y. A. (2000) Transmembrane molecular pump activity of Niemann-Pick C1 protein. Science 290, 2295–2298

99. Sohaebuddin, S. K., and Tang, L. (2013) A simple method to visualize and assess the integrity of lysosomal membrane in mammalian cells using a fluorescent dye. Methods Mol Biol 991, 25–31

100. Bartz, F., Kern, L., Erz, D., Zhu, M., Gilbert, D., Meinhof, T., Wirkner, U., Erfle, H., Muckenthaler, M., Pepperkok, R., and Runz, H. (2009) Identification of cholesterol-regulating genes by targeted RNAi screening. Cell Metab 10, 63–75

101. Terasaki, M., Jaffe, L. A., Hunnicutt, G. R., and Hammer, J. A., 3rd. (1996) Structural change of the endoplasmic reticulum during fertilization: evidence for loss of membrane continuity using the green fluorescent protein. Dev Biol 179, 320–328

102. Qadri, Y. J., Cormet-Boyaka, E., Rooj, A. K., Lee, W., Parpura, V., Fuller, C. M., and Berdiev, B. K. (2012) Low temperature and chemical rescue affect molecular proximity of DeltaF508-cystic fibrosis transmembrane conductance regulator (CFTR) and epithelial sodium channel (ENaC). J Biol Chem 287, 16781–16790

103. Wang, X., Koulov, A. V., Kellner, W. A., Riordan, J. R., and Balch, W. E. (2008) Chemical and biological folding contribute to temperature-sensitive DeltaF508 CFTR trafficking. Traffic 9, 1878–1893

104. Wang, X., Matteson, J., An, Y., Moyer, B., Yoo, J. S., Bannykh, S., Wilson, I. A., Riordan, J. R., and Balch, W. E. (2004) COPII-dependent export of cystic fibrosis transmembrane conductance regulator from the ER uses a di-acidic exit code. J Cell Biol 167, 65–74

105. Liu, B., Li, H., Repa, J. J., Turley, S. D., and Dietschy, J. M. (2008) Genetic variations and treatments that affect the lifespan of the NPC1 mouse. J Lipid Res 49, 663–669

106. Lagace, D. C., and Nachtigal, M. W. (2004) Inhibition of histone deacetylase activity by valproic acid blocks adipogenesis. J Biol Chem 279, 18851–18860

107. Venkataramani, V., Rossner, C., Iffland, L., Schweyer, S., Tamboli, I. Y., Walter, J., Wirths, O., and Bayer, T. A. (2010) Histone deacetylase inhibitor valproic acid inhibits cancer cell proliferation via down-regulation of the alzheimer amyloid precursor protein. J Biol Chem 285, 10678–10689

108. Detich, N., Bovenzi, V., and Szyf, M. (2003) Valproate induces replication-independent active DNA demethylation. J Biol Chem 278, 27586–27592

109. Rauniyar, N., Subramanian, K., Lavallee-Adam, M., Martinez-Bartolome, S., Balch, W. E., and Yates, J. R., 3rd. (2015) Quantitative Proteomics of Human Fibroblasts with I1061T Mutation in Niemann-Pick C1 (NPC1) Protein Provides Insights into the Disease Pathogenesis. Mol Cell Proteomics 14, 1734–1749

110. Subramanian, K., Rauniyar, N., Lavallee-Adam, M., Yates, J. R., 3rd, and Balch, W. E. (2017) Quantitative Analysis of the Proteome Response to the Histone Deacetylase Inhibitor (HDACi) Vorinostat in Niemann-Pick Type C1 disease. Mol Cell Proteomics 16, 1938–1957

111. Arrowsmith, C. H., Bountra, C., Fish, P. V., Lee, K., and Schapira, M. (2012) Epigenetic protein families: a new frontier for drug discovery. Nat Rev Drug Discov 11, 384–400

112. Falkenberg, K. J., and Johnstone, R. W. (2014) Histone deacetylases and their inhibitors in cancer, neurological diseases and immune disorders. Nat Rev Drug Discov 13, 673–691

113. Cornford, E. M., Diep, C. P., and Pardridge, W. M. (1985) Blood-brain barrier transport of valproic acid. Journal of neurochemistry 44, 1541–1550

114. Kakee, A., Takanaga, H., Hosoya, K., Sugiyama, Y., and Terasaki, T. (2002) In vivo evidence for brain-to-blood efflux transport of valproic acid across the blood-brain barrier. Microvascular research 63, 233–238

115. Xuan, A. G., Pan, X. B., Wei, P., Ji, W. D., Zhang, W. J., Liu, J. H., Hong, L. P., Chen, W. L., and Long, D. H. (2015) Valproic acid alleviates memory deficits and attenuates amyloid-beta deposition in transgenic mouse model of Alzheimer’s disease. Mol Neurobiol 51, 300–312

116. Feng, H. L., Leng, Y., Ma, C. H., Zhang, J., Ren, M., and Chuang, D. M. (2008) Combined lithium and valproate treatment delays disease onset, reduces neurological deficits and prolongs survival in an amyotrophic lateral sclerosis mouse model. Neuroscience 155, 567–572

117. Angles, F., Hutt, D. M., and Balch, W. E. (2019) HDAC inhibitors rescue multiple disease-causing CFTR variants. Hum Mol Genet 28, 1982–2000

118. Hutt, D. M., Roth, D. M., Vignaud, H., Cullin, C., and Bouchecareilh, M. (2014) The histone deacetylase inhibitor, Vorinostat, represses hypoxia inducible factor 1 alpha expression through translational inhibition. PLoS One 9, e106224

119. Roth, D. M., Hutt, D. M., Tong, J., Bouchecareilh, M., Wang, N., Seeley, T., Dekkers, J. F., Beekman, J. M., Garza, D., and Drew, L. (2014) Modulation of the maladaptive stress response to manage diseases of protein folding. PLoS biology 12, e1001998

120. de Duve, C., de Barsy, T., Poole, B., Trouet, A., Tulkens, P., and Van Hoof, F. (1974) Commentary. Lysosomotropic agents. Biochem Pharmacol 23, 2495–2531

121. Lu, S., Sung, T., Lin, N., Abraham, R. T., and Jessen, B. A. (2017) Lysosomal adaptation: How cells respond to lysosomotropic compounds. PLoS One 12, e0173771

122. Allfrey, V. G., Faulkner, R., and Mirsky, A. E. (1964) Acetylation and Methylation of Histones and Their Possible Role in the Regulation of Rna Synthesis. Proceedings of the National Academy of Sciences of the United States of America 51, 786–794

123. Kim, S. C., Sprung, R., Chen, Y., Xu, Y., Ball, H., Pei, J., Cheng, T., Kho, Y., Xiao, H., Xiao, L., Grishin, N. V., White, M., Yang, X. J., and Zhao, Y. (2006) Substrate and functional diversity of lysine acetylation revealed by a proteomics survey. Mol Cell 23, 607–618

124. Kovacs, J. J., Murphy, P. J., Gaillard, S., Zhao, X., Wu, J. T., Nicchitta, C. V., Yoshida, M., Toft, D. O., Pratt, W. B., and Yao, T. P. (2005) HDAC6 regulates Hsp90 acetylation and chaperone-dependent activation of glucocorticoid receptor. Mol Cell 18, 601–607

125. Choudhary, C., Weinert, B. T., Nishida, Y., Verdin, E., and Mann, M. (2014) The growing landscape of lysine acetylation links metabolism and cell signalling. Nat Rev Mol Cell Biol 15, 536–550

126. Marinova, Z., Ren, M., Wendland, J. R., Leng, Y., Liang, M. H., Yasuda, S., Leeds, P., and Chuang, D. M. (2009) Valproic acid induces functional heat-shock protein 70 via Class I histone deacetylase inhibition in cortical neurons: a potential role of Sp1 acetylation. Journal of neurochemistry 111, 976–987

127. Leng, Y., Marinova, Z., Reis-Fernandes, M. A., Nau, H., and Chuang, D. M. (2010) Potent neuroprotective effects of novel structural derivatives of valproic acid: potential roles of HDAC inhibition and HSP70 induction. Neuroscience letters 476, 127–132

128. Praggastis, M., Tortelli, B., Zhang, J., Fujiwara, H., Sidhu, R., Chacko, A., Chen, Z., Chung, C., Lieberman, A. P., Sikora, J., Davidson, C., Walkley, S. U., Pipalia, N. H., Maxfield, F. R., Schaffer, J. E., and Ory, D. S. (2015) A murine Niemann-Pick C1 I1061T knock-in model recapitulates the pathological features of the most prevalent human disease allele. J Neurosci 35, 8091–8106

129. Chittur, S. V., Sangster-Guity, N., and McCormick, P. J. (2008) Histone deacetylase inhibitors: a new mode for inhibition of cholesterol metabolism. BMC genomics 9, 507

130. Munkacsi, A. B., Hammond, N., Schneider, R. T., Senanayake, D. S., Higaki, K., Lagutin, K., Bloor, S. J., Ory, D. S., Maue, R. A., Chen, F. W., Hernandez-Ono, A., Dahlson, N., Repa, J. J., Ginsberg, H. N., Ioannou, Y. A., and Sturley, S. L. (2017) Normalization of Hepatic Homeostasis in the Npc1(nmf164) Mouse Model of Niemann-Pick Type C Disease Treated with the Histone Deacetylase Inhibitor Vorinostat. J Biol Chem 292, 4395–4410

131. Nunes, M. J., Moutinho, M., Gama, M. J., Rodrigues, C. M., and Rodrigues, E. (2013) Histone deacetylase inhibition decreases cholesterol levels in neuronal cells by modulating key genes in cholesterol synthesis, uptake and efflux. PLoS One 8, e53394

132. Taylor, A. M., Liu, B., Mari, Y., Liu, B., and Repa, J. J. (2012) Cyclodextrin mediates rapid changes in lipid balance in Npc1-/-mice without carrying cholesterol through the bloodstream. J Lipid Res 53, 2331–2342

133. Millard, E. E., Gale, S. E., Dudley, N., Zhang, J., Schaffer, J. E., and Ory, D. S. (2005) The sterol-sensing domain of the Niemann-Pick C1 (NPC1) protein regulates trafficking of low density lipoprotein cholesterol. J Biol Chem 280, 28581–28590

134. Tangemo, C., Weber, D., Theiss, S., Mengel, E., and Runz, H. (2011) Niemann-Pick Type C disease: characterizing lipid levels in patients with variant lysosomal cholesterol storage. J Lipid Res 52, 813–825

135. Hao, M., Lin, S. X., Karylowski, O. J., Wustner, D., McGraw, T. E., and Maxfield, F. R. (2002) Vesicular and non-vesicular sterol transport in living cells. The endocytic recycling compartment is a major sterol storage organelle. J Biol Chem 277, 609–617

