## Supplemental Figures for "Correction of Niemann-Pick type C1 disease with the histone deacetylase inhibitor valproic acid"

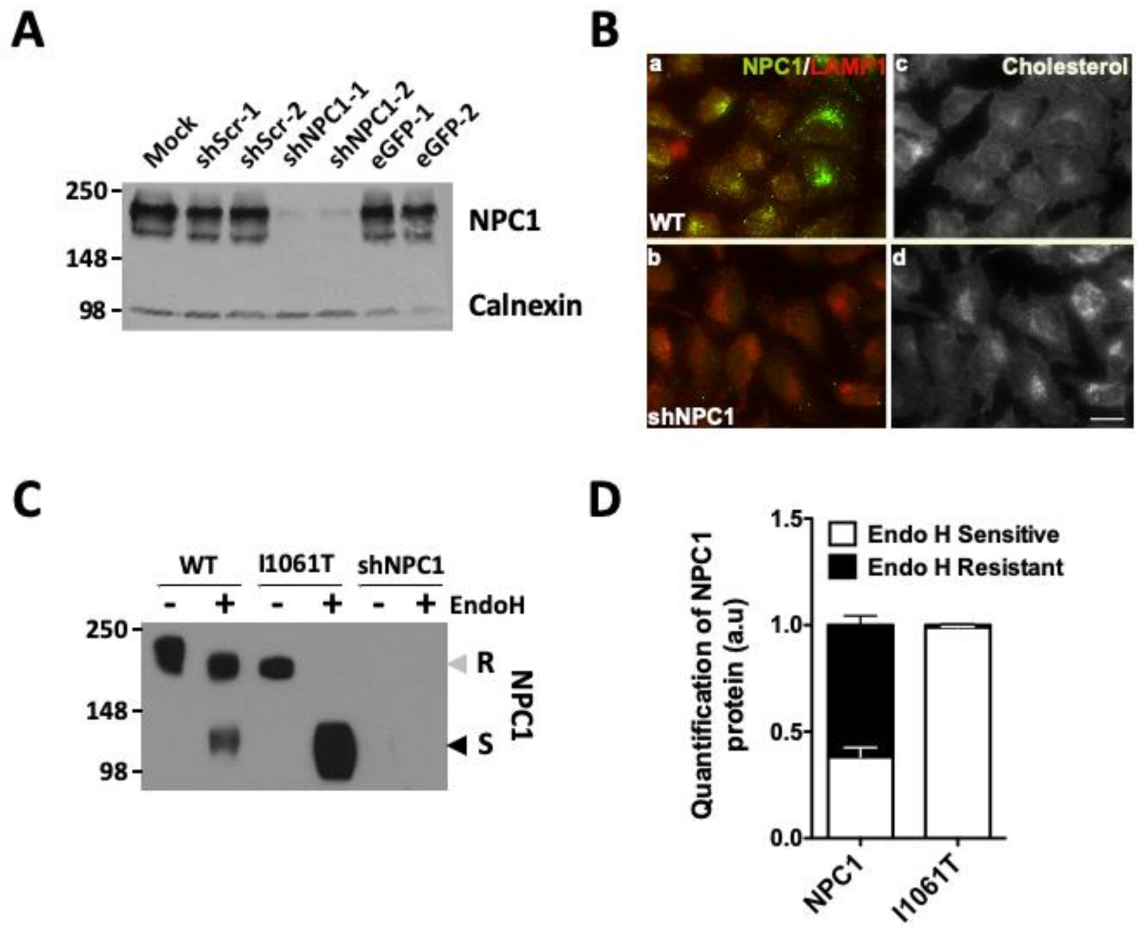


**Figure S1. Development of HeLa cells stably expressing NPC1 variants. A.** Western blot analysis of NPC1 from HeLa cells stably transduced with lentiviral particles delivering the indicated plasmid. **B.** Representative immunofluorescent images of control and shNPC1 transduced HeLa cells. Shown are NPC1 (green) and LAMP1 (red) staining (panels a-b) as well as filipin staining (panels c-d). The scale bar shown represents 10 µm. **C.** Representative Western blot of NPC1 from control HeLa-shNPC1 cells and HeLa-shNPC1 stably transduced with WT- or I1061T-NPC1. Lysates were treated without (-) or with (+) endoH prior to SDS-PAGE. **D.** Bar graph depicting the amount of EndoH$^{S}$ and EndoH$^{R}$ glycoforms as a fraction of total NPC1 from HeLa-shNPC1 cells stably expressing WT- or I1061T-NPC1. The data shown represents mean ± SD (n = 3).

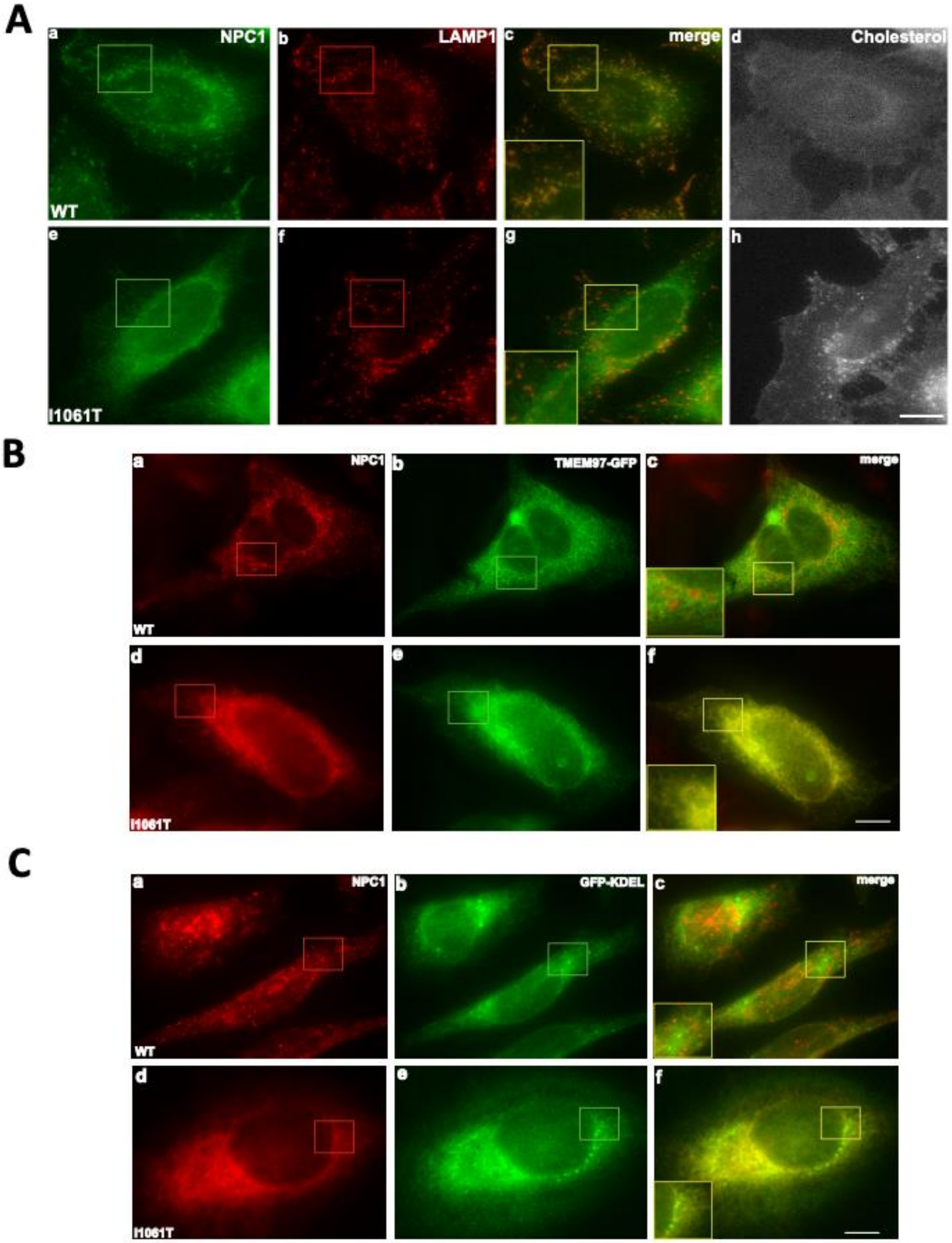


**Figure S2. Mislocalization of human NPC1 I1061T in HeLa-shNPC1 cells. A.** Representative immunofluorescent images of HeLa-shNPC1 cells stably expressing human WT- (Top) and I1061T-NPC1 (Bottom). Shown are NPC1 (green) (panels a & e), LAMP1 (red) (panels b & f) and filipin (greyscale) (panels d & h) staining. **B.** Representative immunofluorescent images of HeLa-shNPC1 cells stably expressing human WT- (Top) and I1061T-NPC1 (Bottom). Shown are NPC1 (red) (panels a & d), TMEM97-GFP (green) (panels b & e). **C.** Representative immunofluorescent images of HeLa-shNPC1 cells stably expressing human WT- (Top) and I1061T-NPC1 (Bottom). Shown are NPC1 (red) (panels a & d), GFP-KDEL (green) (panels b & e). In all panels the scale bar shown represents 10 μm.

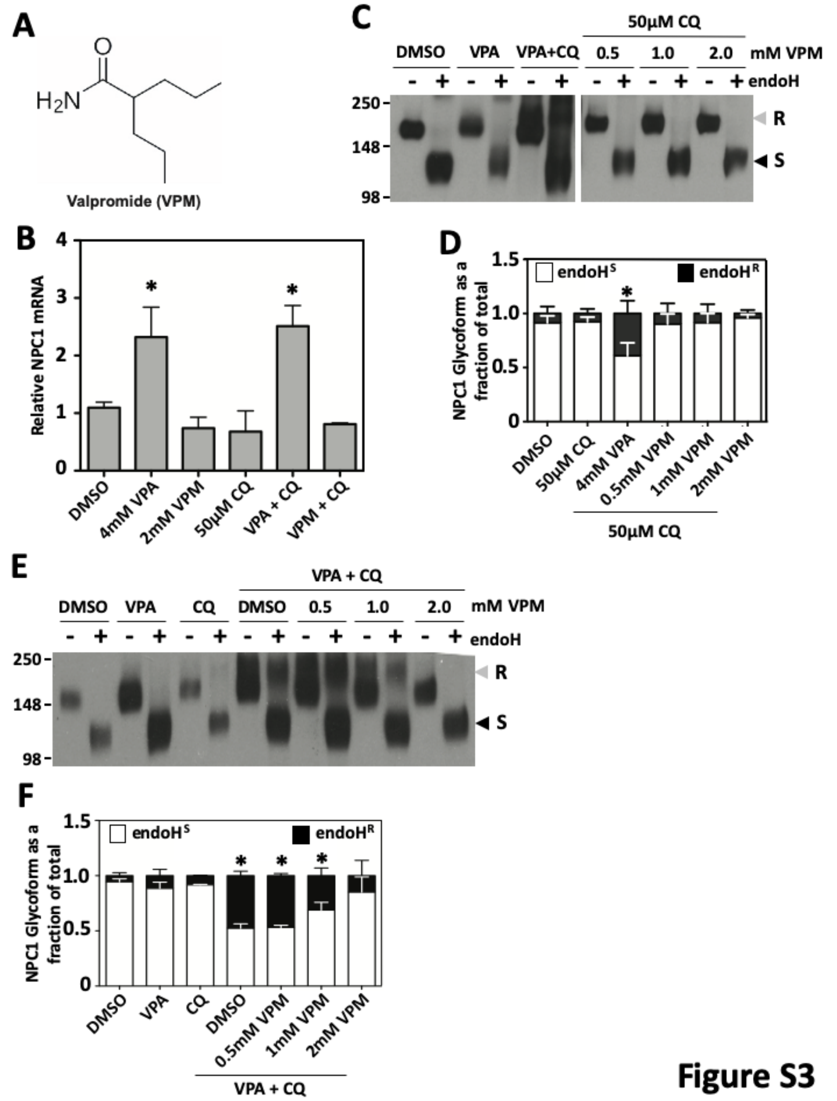


Figure S3

**Figure S3. Valpromide (VPM) does not correct the trafficking of I1061T-NPC1. A.** Chemical structure of valpromide. **B.** Bar graph depicting the NPC1 mRNA levels in I1061T-expressing HeLa-shNPC1 cell lysates treated with DMSO, 4mM VPA, 2mM VPM, 50μM CQ or a combination of these compounds for 24 h. The data is shown as the relative mean ± SD with the vehicle treatment normalized to 1 and asterisks indicates $p < 0.05$ as determined by a one-way ANOVA and Dunnett's post-tests (n = 3). **C.** Representative Western blot of I1061T-expressing HeLa-shNPC1 cell lysates treated with DMSO, 4mM VPA, 4mM VPA + 50μM CQ or the indicated concentrations of VPM in the presence of 50μM CQ for 24 h. NPC1 was immunoprecipitated and treated without (-) or with (+) Endo H prior to SDS-PAGE. **D.** Bar graph depicting the amount of EndoH$^S$ (white) and EndoH$^R$ (black) glycoforms as a fraction of total NPC1 from I1061T-expressing HeLa-shNPC1 cell lysates treated as in panel **C**. The data are presented as the normalized mean ± SD and the asterisk indicates $p < 0.05$ using a two-tailed T-test with DMSO treatment as a reference (n = 3). **E.** Representative Western blot of I1061T-expressing HeLa-shNPC1 cell lysates treated with DMSO, 4mM VPA, 50μM CQ or a combination of these treatments in the presence of the indicated concentrations of VPM for 24 h. NPC1 was immunoprecipitated and treated without (-) or with (+) Endo H prior to SDS-PAGE. **F.** Bar graph depicting amount of EndoH$^S$ (white) and EndoH$^R$ (black) glycoforms as a fraction of total NPC1 from I1061T-expressing HeLa-shNPC1 cell lysates treated as in panel **E**. The data are presented as the normalized mean ± SD and the asterisk indicates $p < 0.05$ using a two-tailed T-test with DMSO treatment as a reference (n = 3).
